# Occludin Acts as a Dynein Adaptor Regulating Permeability and Collateral Angiogenesis

**DOI:** 10.1101/2025.06.12.659326

**Authors:** Xuwen Liu, Enming J. Su, Alyssa Dreffs, John P. Gillies, Madeline Merlino, Lu Gao, Julian S. Peregoff, Cheng-mao Lin, Margaret Elizabeth Ross, Morgan E. DeSantis, Daniel A. Lawrence, David A. Antonetti

## Abstract

Previous studies of the tight junction protein occludin (OCLN) suggest that multiple phosphorylation sites on the carboxy-terminal domain contribute a regulatory role in vascular barrier properties. However, gene deletion studies failed to identify a clear functional role for OCLN, despite multiple phenotypic alterations. Importantly, previous studies targeting exon 3 allowed expression of a splice variant starting at exon 4, (isoform 4) that expresses the full carboxy-terminal tail. Here we show that the OCLN carboxy-terminus forms a complex with the light intermediate chain (LIC) of dynein to link tight junction cargo to the minus end directed motor protein. Mutational analysis revealed S471 phosphorylation is required for binding to the LIC while S490 phosphorylation is required for trafficking. Expressing *OCLN* S490A mutant prevented endothelial cell proliferation and collateral angiogenesis. *OCLN* gene deletion targeting exon 5, preventing full length and isoform 4 expression, resulted in embryonic lethality. In summary, OCLN links tight junction cargo to the dynein motor, regulating trafficking in a phosphorylation dependent manner and contributing to both VEGF-induced vascular permeability and collateral angiogenesis.

## Introduction

Occludin (OCLN) was the first transmembrane tight junction protein identified ^1^; however, the function of occludin has remained elusive. OCLN has sequence homology to MARVEL (<u>m</u>yelin/lymphocyte (MAL) <u>a</u>nd related proteins for <u>ve</u>sicle trafficking and membrane link) proteins with conserved sequence adjacent to the four-transmembrane regions that are characterized by their involvement in vesicle trafficking ^2^. OCLN is now also recognized as among the TAMP or tight junction <u>a</u>ssociated <u>M</u>ARVEL <u>p</u>rotein family including MARVELD1 or OCLN, MARVELD2 or tricellulin and MARVELD3. OCLN (522aa) consists of a 66aa N-terminal cytoplasmic domain, transmembrane (TM)1 (23aa), extracellular loop (EL)1 (46aa), TM2 (25aa), intracellular loop (10aa), TM3 (25aa), EL2 (48aa), TM4 (22aa) and a long carboxy terminus of 257aa ^3^. The carboxy terminus includes a coiled coil domain (CC residues 413-522) that binds to ZO-1 ^4^, 2 ^5^ and 3 ^6^. This carboxy terminus is heavily phosphorylated in the CC domain and adjacent to the CC (reviewed in ^3,7^).

OCLN phosphorylation has been shown to regulate barrier properties. There is evidence for Tyr phosphorylation regulating interaction with the tight junction organizing protein zonula occludens 1 (ZO-1) ^8^ and Ser/Thr phosphorylation regulating epithelial junction integrity ^9^ assembly ^10^, and paracellular permeability through dynamic changes in tight junction protein interactions ^11^. Multiple phosphorylation sites were identified in vascular endothelial cells through mass spectrometry analysis ^12^, two of which are in the CC domain and have been found to have functional significance demonstrated by mutational analysis. Phosphorylation of the S471 site in OCLN is required for post-confluence, size reductive proliferation and for maturation of epithelial cell junctional complex ^13^. Phosphorylation of S490 by PKCβ is stimulated by vascular endothelial growth factor (VEGF) in endothelial cells and regulates permeability ^14,15^. Expression of S490A acts in a dominant manner to block VEGF-induced permeability in endothelial cells ^14,15^ and expression of this mutant in transgenic animals inhibits VEGF- or diabetes-induced retinal vascular permeability preserving visual function ^16^ and reduces stroke induced-permeability and allows late tissue plasminogen activator treatment of stroke without hemorrhagic transformation ^17^.

Genetic evidence has clearly demonstrated a role for the claudin proteins in barrier properties. Claudin proteins are a large family of tetraspan tight junction proteins of at least 24 different genes in humans that confer barrier properties dependent on which isoforms are expressed with some claudins acting in barrier formation and others as charge and size selective paracellular pores ^18–20^. Claudin 5 expression is restricted to vascular endothelial cells and has been the only claudin shown to confer barrier properties to the blood-brain barrier through transgenic studies as claudin 5 gene deletion leads to death within 10h after birth associated with flux of molecules <800Da into the brain ^21,22^.

Studies of gene deletion of *Ocln* have been less informative regarding the role of this gene product. Gene deletion through targeting exon 3 with flanking lox (flox) sites revealed mice with a host of phenotypic alterations such as brain calcification, male infertility, and gut hyperplasia of the gastric epithelium, thinning of compact bone and male infertility but tight junctions and barrier properties appeared intact ^23,24^. However, more recent data reveals that these *Ocln*^fl3/fl3^ transgenic mice allowed expression of a splice variant from an internal ribosome entry site at exon 4 called ΔN or isoform 4 (Iso4) OCLN that includes 14 amino acids in the transmembrane region and the carboxy-terminal tail including phosphorylation sites S471 and S490 ^25^.

Multiple studies have demonstrated that OCLN contributes to cell proliferation in many cell types. OCLN, phosphorylated at S490, was localized to cell centrosomes and shown to be necessary for VEGF driven EC proliferation in culture ^26^, and in epithelial cells the S490A mutation delays mitotic entry ^27^. Phosphorylation of the S471 site in OCLN is required for post-confluence, size reductive proliferation and for maturation of epithelial cell junctional complex ^13^. Further, OCLN^fl3/fl3^ deletion contributes to gastric epithelium hyperplasia ^24^ and OCLN was identified as an anti-oncogene ^28^. In humans, various OCLN mutations lead to microcephaly, brain calcification and renal disease ^29–31^. Analysis of cortical neuroprogenitor cell proliferation reveals OCLN localizes to centrosomes and deletion of full-length OCLN leads to alterations in mitotic spindle pole organization reducing progenitor self-renewal and increasing apoptosis in both mice and human brain organoids ^25^. Together, these studies provide compelling evidence that OCLN contributes critical activities regulating cell growth and proliferation, specifically in cells that possess junctional complexes and polarization.

In the current study, we provide evidence that the OCLN carboxy terminus acts to link the tight junction complex to the microtubule minus end directed motor dynein, allowing internalization of the junctional complex and paracellular permeability. We find that OCLN also traffics to the centrosome to support cell proliferation. Expression of S490A *OCLN* in mice prevents post-ischemia collateral vessel growth in brain and retina. *OCLN* mutational analysis reveals the carboxy terminus is necessary for trafficking to centrosomes in a S490 phosphorylation dependent manner. Further the C-terminus of OCLN co-precipitates with the light intermediate chain (LIC) of dynein. S490A Iso4 of OCLN can act in a dominant negative manner blocking VEGF-induced permeability and proliferation of endothelial cells like full length S490A OCLN. Finally, a new gene deletion model targeting exon 5 (OCLN^fl5/fl5^), that ablates expression of both full length and Iso4 OCLN, is embryonic lethal associated with blood vessel rupture and hemorrhaging with some animals demonstrating gastroschisis.

## Results

### Expression of *OCLN*S490A impedes collateral vessel growth

We previously showed that expression of *OCLN*S490A mutant in endothelial cells prevented diabetes induced retinal vascular permeability ^16^ and stroke induced permeability in the brain and prevented late tPA induced hemorrhagic transformation ^17^ clearly demonstrating a role for this phosphorylation of OCLN in regulation of vascular permeability. To determine whether OCLNS490 phosphorylation was required for collateral vessel angiogenesis in the transgenic animals, we developed a laser induced model of branched retinal vein occlusion (BRVO). Retinal veins were ablated by a brief pulse from a 532nm laser using a Micron III retinal imaging system and collateral vessel growth was monitored in retinal whole mounts at various times after injury. We examined vessel growth in *OCLN*S490A^+/+^ mice compared to WT*OCLN* with expression controlled by a floxed stop site after the CAG promoter (described in ^16^). Inducible PDGFiCre was used for vascular endothelial specific expression. Mice were *Ocln*^fl3/fl3^ with deletion of endogenous full length *Ocln* but maintained Iso4 *Ocln* after Cre excision and simultaneous expression of S490A or WT transgene. Immunofluorescence staining of the retinal whole mount in littermate control (LM ctrl) animals at day 7 after BRVO revealed dramatic increase in cell proliferation indicated by Ki-67 positive staining, some co-labeled with CD31 while others were in the retinal parenchyma (Fig. 1a), clearly indicating both endothelial and non-endothelial proliferation. The number of Ki-67 positive cells in superficial plexus was significantly increased in LM Ctrl, Cre+, and WT*OCLN* groups after BRVO as compared to B6 Ctrl non-laser group (Fig. 1b), while the increased number of Ki-67 positive cells was significantly inhibited in S490A group after BRVO (Fig. 1b and c). The result indicates that endothelial restricted expression of transgenic *OCLN*WT had no effect on this proliferation but endothelial expression of *OCLN*S490A mutant prevented the increase in cell proliferation of both endothelial and non-endothelial cells in BRVO. Previous studies suggested the time frame of 2-4 weeks for retinal collateral vessel growth ^32,33^. Therefore, we evaluated the role of *OCLN*S490A on collateral vessels growth at 4 weeks after BRVO. Immunofluorescence staining of retinal whole mounts with CD31 antibody revealed there were more vessels surrounding the occluded vein in superficial plexus in LM ctrl after BRVO as compared to its non-laser group and *OCLN*S490A inhibited this vessel proliferation (Fig. 1d). Quantification analysis demonstrated that expression of *OCLN*S490A mutant completely prevented the BRVO induced increase in capillary collateral vessel area and length observed in the Cre-littermate controls (LM Ctrl) (Fig. 1e and f). The proliferation experiments were repeated with endogenous *Ocln* background, (i.e. without *Ocln* exon 3 excision). Again, expression of *OCLN*S490A prevented BRVO-induced cell proliferation almost identical to the studies with exon 3 excision (Fig. 1g and h), indicating the dominant effect of *OCLN*S490A in the presence of full length and Iso4 *Ocln*. We also explored the laser-induced choroidal neovascularization (CNV) angiogenesis mouse model and observed a modest but not significant loss of vessel growth with *OCLN*S490A expression in these vessels (Supplementary Fig.1a and 1b). It is noteworthy these vessels have limited tight junction expression compared to retinal vessels.

**Figure 1.**
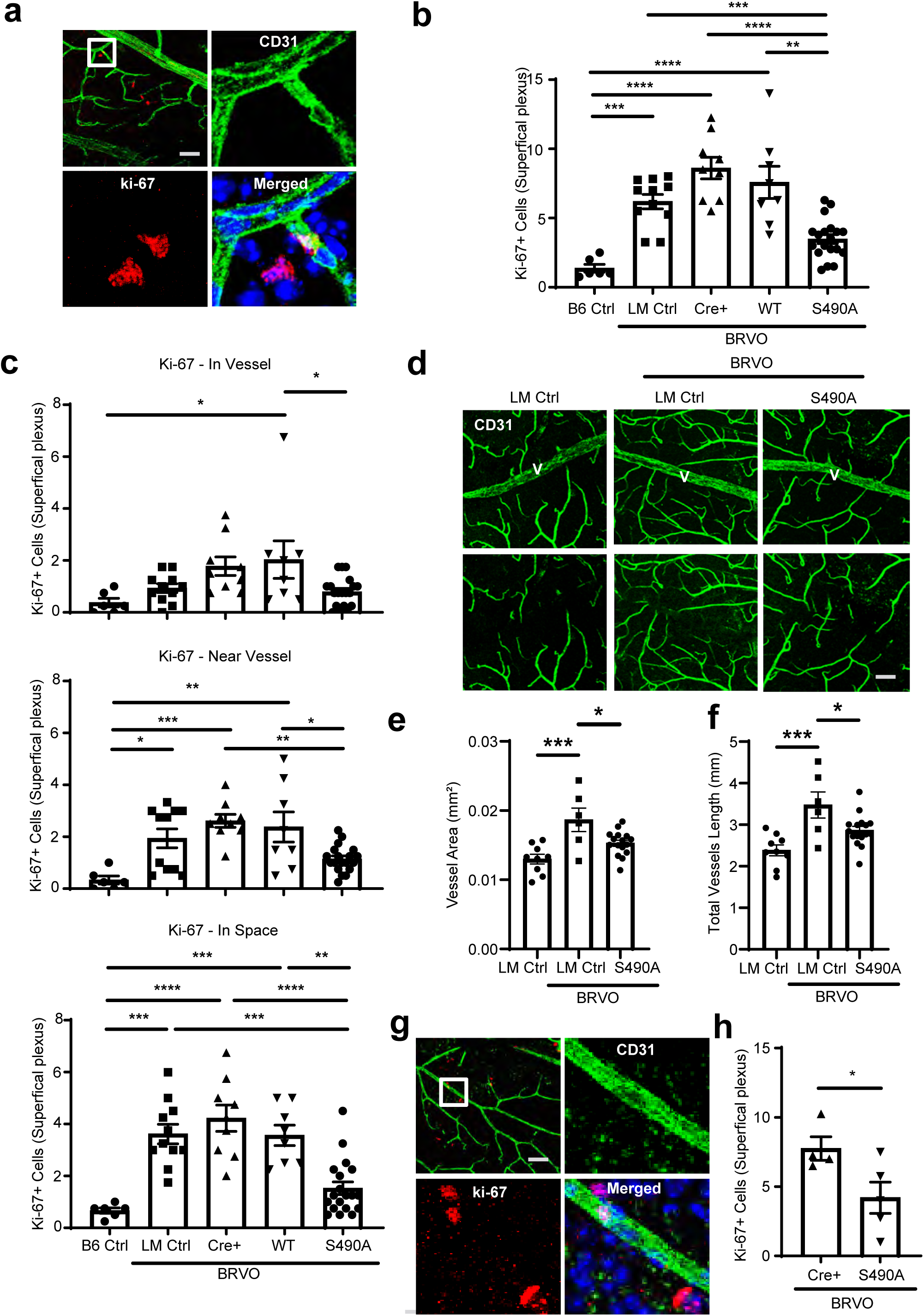
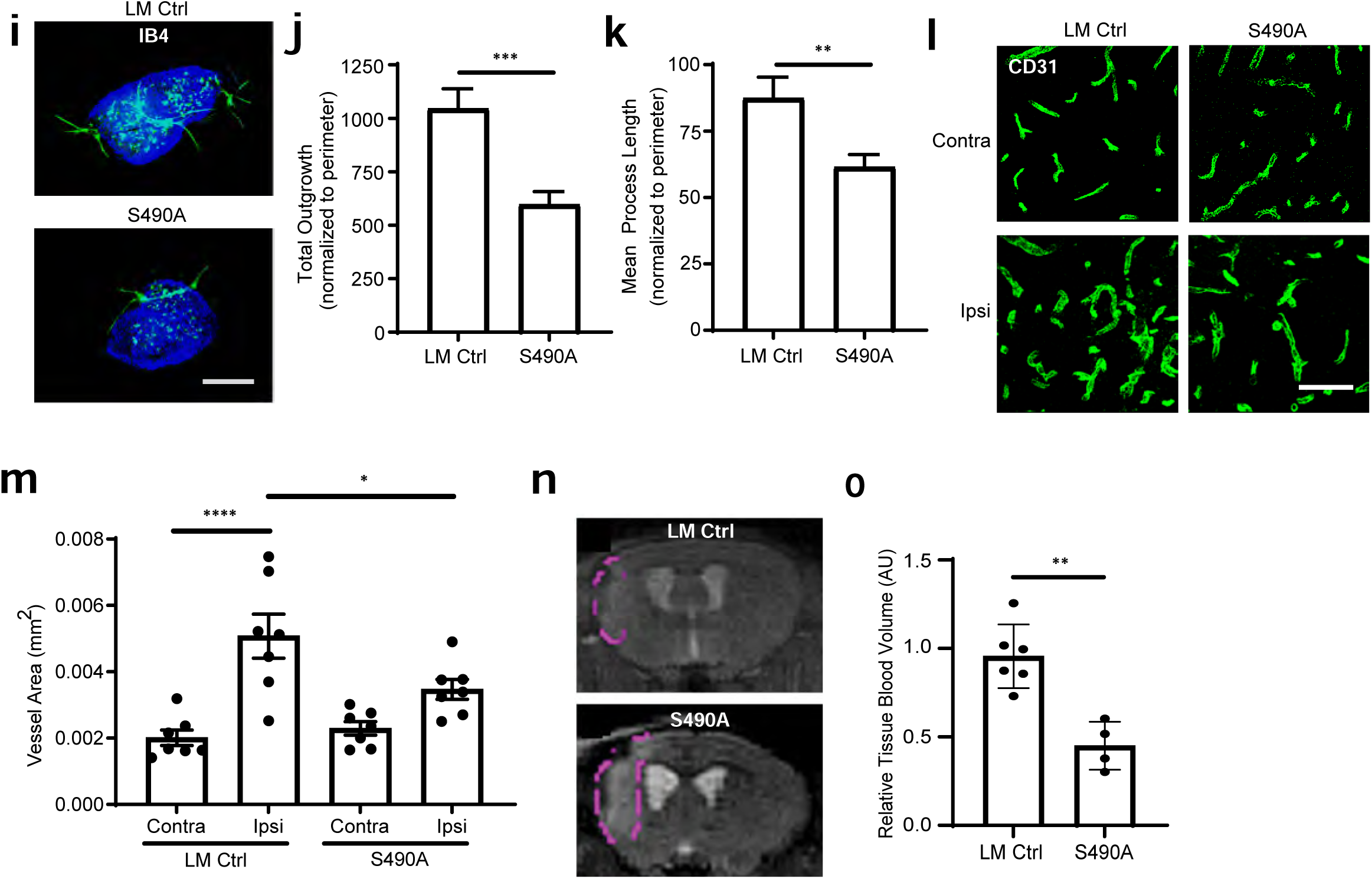
Expression of occludin S490A impedes collateral angiogenesis. **a.** Whole mount retinal images of CD31 and Ki-67 immunostaining. Scale bar: 50 µm. **(b** and **c)** Quantification of Ki-67 positive cells. Mouse lines; B6 Ctrl: C57BL/6J; littermate (LM) Ctrl: *Pdgfb-iCre*ER^-^, *Ocln*^fl3/fl3^, *OCLN*S490A^+/+^; Cre^+^: *Pdgfb-iCre*ER^+^, *Ocln*^fl3/fl3^; WT: *Pdgfb-iCre*ER^+^, *Occ*^fl3/fl3^, WT*OCLN*^+/+^; S490A: *Pdgfb-iCre*ER^+^, *Ocln*^fl3/fl3^, *OCLN*S490A^+/+^. **(d)** Confocal images of CD31 stained vessels at 4 weeks after BRVO showing targeted vein (V) and image after removal of V using Photoshop (Adobe). Scale bars: 50 µm. **(e-f)** Quantitation of vessels using Angi Tool. Expression of *OCLN* S490A reduced endothelial cell proliferation in mice carrying endogenous *Ocln* at 1 week after BRVO. **(g)** Representative images, scale bar: 50 µm and **(h)** quantification. Mouse lines: Cre^+^: *Tek-Cre*^+^, *OCLN*S490A^-^/^-^; S490A: *Tek-Cre*^+^, *OCLN*S490A^+^/^+^. **(i)** Retinal explants with IB4 (green) and Hoechst (blue) immunostaining. A total of 16 retinal explants culture was established for each eye, 6 eyes in each group. **(j-k)** Quantification by MetaMorph software. LM Ctrl: *Pdgfb-iCreER*^-^, *Ocln*^fl3/fl3^, *OCLN*S490A^+/+^; S490A: *Pdgfb*-*iCreER*^+^, *Ocln*^fl3/fl3^, *OCLN*S490A^+/+^. **(l-m)** Neovascularization was inhibited in S490A mice at 3 weeks after MCAO. **(l)** Images of CD31 stained vessels in peri-infarct areas at three weeks after stroke. Four regions in each section were quantified and three sections per mouse were evaluated (12 images per mouse, n=1). Scale bar: 50 µm. **(m)** Quantitative analysis by Angi Tool image software. LM Ctrl: *Pdgfb-iCreER*^-^, *Ocln*^fl3/fl3^, *OCLN*S490A^+/+^; S490A: *Pdgfb-iCreER*^+^, *Ocln*^fl3/fl3^, *OCLN*S490A^+/+^. Contra: Contralateral region, Ipsi: Ipsilateral region. **(n)** MRI imaging of cerebral blood volume. LM Ctrl and S490A mice underwent MCAO and 21 days later, T2-weighted MRI images were obtained from before gadolinium injection through 30min after injection. **(o)** Regions of interest from the infarct zone were selected and computed for cerebral blood volume. Mouse lines are the same as **l** and **m**.

The effect of *OCLN*S490A expression on vessel growth was supported by an *ex vivo* retinal explant experiment. As shown in Fig. 1i, vascular outgrowth from retinal fragments was significantly decreased from *OCLN*S490A mice compared to LM controls. Quantification of IB4 stained vessels revealed a decrease in total vessel outgrowth and mean process length with *OCLN*S490A expression (Fig. 1j and k). These studies reveal the inhibitory effect does not require circulating endothelial precursors as the inhibitory effect of *OCLN*S490A expression on angiogenesis was observed in the *ex vivo* model.

Lastly, we tested if *OCLN*S490A had similar inhibitory effect on angiogenesis in the brain after stroke. Stroke was induced by photothrombotic MCAO and immunofluorescence staining of brain cross sections with CD31 antibody was completed followed by vessel area quantification. In LM control animals MCAO induced a 2.5-fold increase in vessel area in the peri-infarct region at 21 days after stroke compared to the contralateral region. However, the increased collateral angiogenesis was significantly reduced in *OCLN*S490A expressing mice (Fig. 1l and m). To corroborate this finding the relative tissue blood volume, as a measure of blood circulation, in the infarct region was measured after MCAO by MRI imaging (Fig. 1n). LM control mice had significantly more tissue blood volume in the infarcted area at 3 weeks after MCAO compared to the animals expressing *OCLN*S490A in the vascular endothelium (Fig. 1o). Taken together, these data provide compelling evidence that OCLN phosphorylation on Ser490 contributes to endothelial cell growth and collateral angiogenesis in the CNS as expression of the phospho-inhibitory *OCLN*S490A inhibits angiogenesis in vessels with well-developed tight junctions.

### OCLN localizes at centrosomes in a membrane vesicle requiring Iso4 and Ser490 phosphorylation

Our previous studies have demonstrated that S490 phosphorylated OCLN localizes to centrosomes throughout mitosis and S490 phosphorylation is required for mitotic entry in epithelial cells ^34^ and VEGF-induced retinal neovascularization ^26^. Here, OCLN centrosomal localization was further explored in U2OS cells by mutational analysis to identify the regions necessary and sufficient for targeting to centrosomes. The U2OS cell line has large and clearly observable centrosomes aiding analysis. First, we performed immunofluorescence staining with centrosome marker γ-tubulin antibody and antibodies against OCLN N-terminus, C-terminus and p490 with all 3 showing clear centrosome co-localization (Fig. 2a). Next, we made a series of epitope-tagged OCLN mutants to determine which domains are necessary for centrosomal localization of OCLN (Fig. 2b). U2OS cells were transfected with OCLN mutants with either N-terminal GFP or C-terminal V5 tags and stained for the tag along with centrosomal marker γ-tubulin and DNA Hoechst dye (Fig. 2c and 2d and quantification 2e and 2f). Both N-terminal GFP tagged (GFP-WT) or C-terminal tagged V5 (WT-V5) forms of OCLN localize to centrosomes. Mutants of the N-terminus through the first and second extracellular loop had no effect on centrosomal localization (Fig. 2e). However, OCLN centrosomal localization was completely blocked by S490A mutation (GFP-S490A) and in the mutants lacking C-terminal coiled-coil domain (GFP-ΔCC). The C-terminal CC domain (GFP-CC) alone failed to lead to centrosomal co-localization (2f). The full C-terminal tail, either as N-terminal GFP-tagged or C-terminal V5-tagged, was also not sufficient to yield centrosomal co-localization (Fig. 2f and g). However, when we created a V5 tagged mimic of the alternatively spliced Iso4 OCLN that possesses the full C-terminal tail plus 14 transmembrane amino acids, we observed centrosomal co-localization like V5 WT-OCLN. Finally, we added a myristylation motif to the C-terminal mutant (Myr-Cter-V5) to provide a membrane anchor. The result indicated that Myr-Cter-V5 restored OCLN centrosomal localization to near 50% (Fig. 2g). Collectively, these studies demonstrate that OCLN centrosomal localization requires membrane association, C-terminal coiled-coil domain, and Ser490 phosphorylation. Further, the Iso4 splice variant, which contains all these features, traffics robustly to centrosomes.

**Figure 2.**
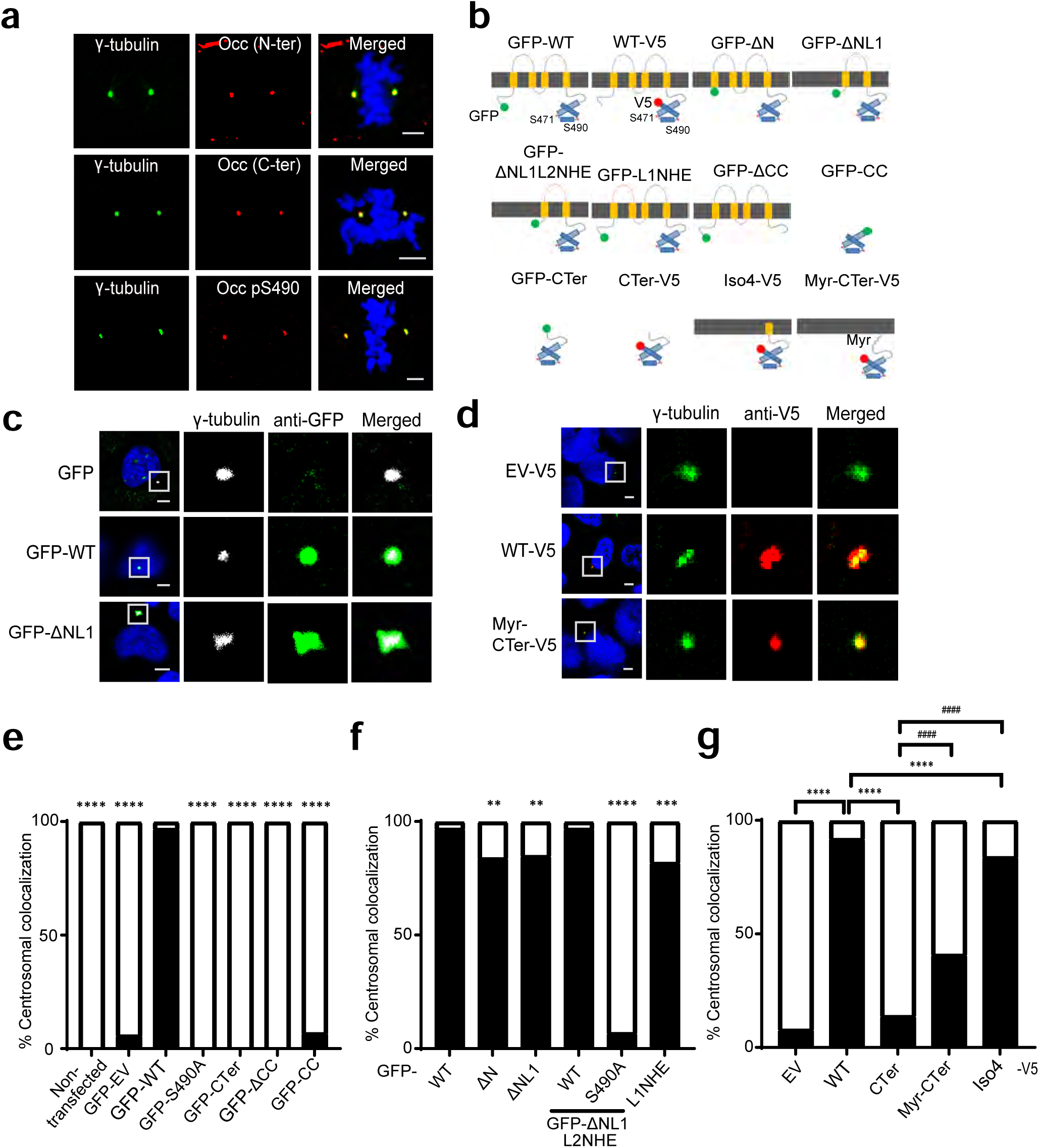
Occludin localization at centrosomes requires membrane component and C-terminal coiled-coil including Ser490 phosphorylation. **(a)** Occludin can be identified at centrosomes of dividing cells by multiple occludin antibodies. Representative confocal images of U2OS cells immunostained with rabbit anti-N-terminal OCLN antibody, mouse anti-C-terminal OCLN antibody, or rabbit anti-OCLN pS490 antibody (all in red) with centrosomal marker γ-tubulin antibody (green) and Hoechst staining for DNA (blue). Scale bar: 5 µm. **(b)** Schematic diagram of GFP or V5 epitope-tagged full-length OCLN and mutants. **(c** and **d)** U2OS cells transfected with *OCLN* mutants in either N-terminal tagged GFP vector (GFP) or C-terminal tagged V5 vector (EV-V5) were fixed and stained with antibodies for GFP (green) and centrosomal marker γ-tubulin (white) or V5 (red) and γ-tubulin (green) mAb. Scale bars: 5 µm. **c** and **d** are the representative images. **(e-g)** Quantification of percent of transfected cells demonstrating co-localization with γ-tubulin at centrosomes. Minimum 30 cells of each condition were analyzed by Chi-Square analysis. Black bars, complete colocalization; open bars, no colocalization. **(e)** Compared to GFP-WT, **P <0.01, ***P<0.001 and ****P<0.0001. **(f)** compared to GFP-WT ****P<0.0001. **(g)** Compared to WT-V5 ****P<0.0001, compared to C-Ter-V5 ^####^P<0.0001

### OCLN Iso4 and Ser490 phosphorylation regulate VEGF-driven proliferation

We next determined whether the dominant Ser490A mutation was effective in controlling VEGF-driven proliferation in Iso4 OCLN as we previously observed for full length OCLN ^26^. BREC were transfected with plasmids containing *OCLN* mutants and compared to cells with wild-type *OCLN* or empty vector. Measures of endothelial proliferation in response to VEGF were carried out by Click-iT EdU DNA synthesis assay in 3D collagen culture. This analysis revealed that transfection of *OCLN*S490A inhibited VEGF-induced DNA synthesis as reported previously ^26^ (Fig. 3a). In addition, VEGF-induced proliferation was inhibited in S490A Iso4 mutant (Iso4-V5 S490A) (Fig. 3b). *OCLN* mutants lacking extracellular loop 1 (GFP-ΔNL1) or with additional mutations to loop2 (GFP-ΔNL1L2NHE and GFP-ΔNL1L2NHE S490A) all effectively blocked VEGF-induced proliferation (Fig. 3c). As both GFP-S490A and GFP-ΔNL1 inhibited VEGF-induced DNA synthesis but OCLN was localized to centrosomes in GFP-ΔNL1 but not GFP-S490A transfected cells, cell cycle analysis in 3D collagen culture in the presence of VEGF was performed to determine if these OCLN mutants effected cell cycle progression. Cell cycle analysis demonstrated that *OCLN* GFP-ΔNL1 expression shortened prophase and prometaphase and prolonged metaphase to telophase as compared to GFP-EV, GFP-WT or GFP-S490A *OCLN* (Fig. 3d). But expression of *OCLN*S490A did not alter cell cycle progression. Further apoptosis assay showed that GFP-ΔNL1 mutation induced apoptosis as compared to GFP control, or WT*OCLN* (Fig. 3e). S490A mutant had a more modest increase in apoptosis compared to GFP-control. Finally, we demonstrated that the Iso4-S490A mutant also blocked VEGF induced permeability in primary endothelial cells as previously reported for full length *OCLN*S490A ^14^ (Fig. 3f). Taken together, these studies demonstrate that OCLN Iso4 with a Ser490A mutation is sufficient to block both VEGF-induced endothelial cell permeability and proliferation and that Iso4 is necessary and sufficient to traffic to centrosomes. Meanwhile, deletions of the N-terminus of OCLN affecting loop1, impacts cell division and lead to apoptosis consistent with the full length *Ocln* gene deletion and impaired neural progenitor cell division previously reported ^25^.

**Figure 3.**
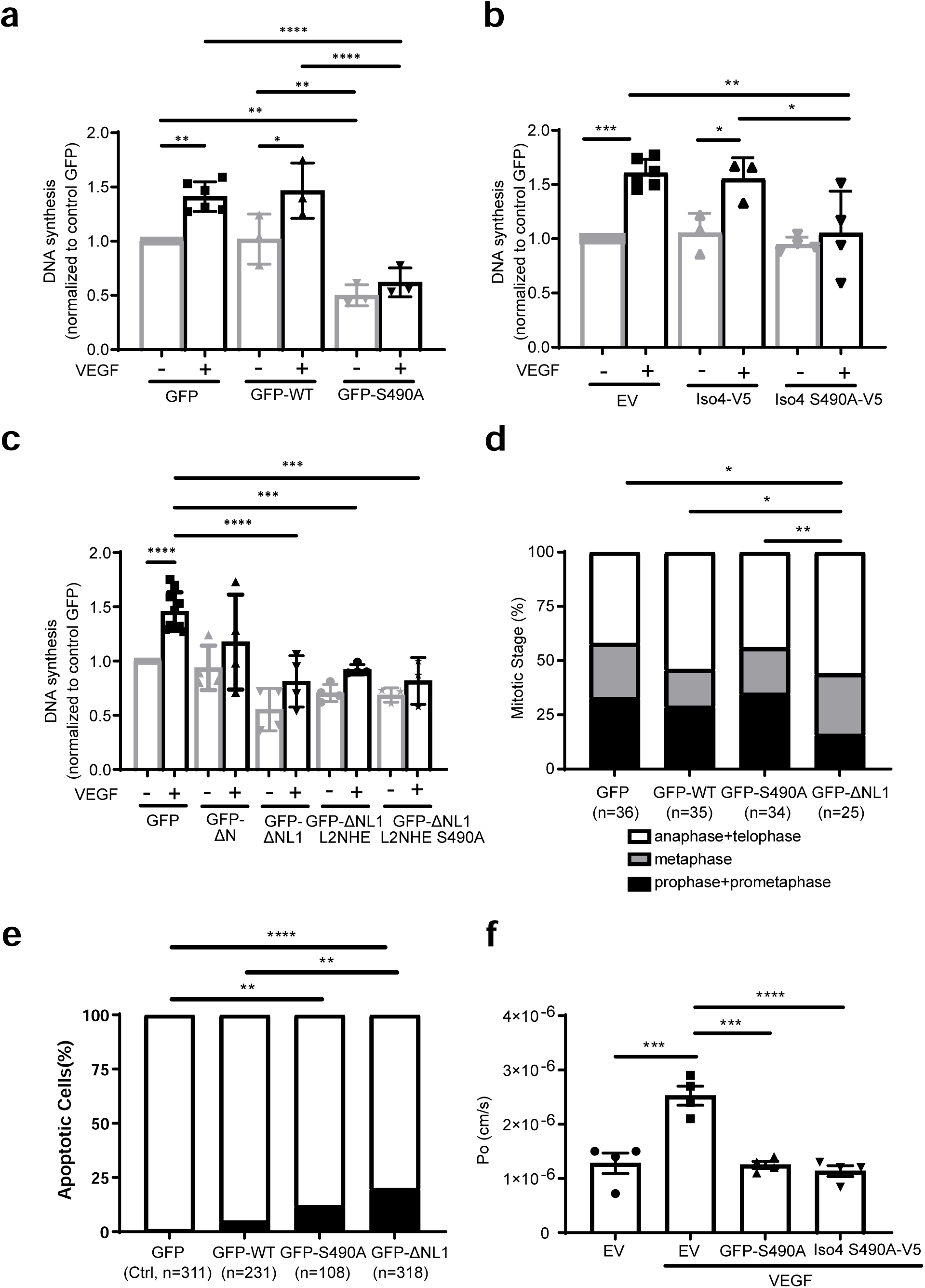
Endothelial cell division and permeability are regulated by occludin Ser490 phosphorylation. **(a-c)** BREC were transfected with GFP, WT-*OCLN* (GFP-WT) or mutants and DNA synthesis was measured by Click-iT EdU at 8 hours after VEGF treatment. Quantification from at least three independent experiments with each experiment from 6 individual wells in a 24-well plate yielding n of 1. **(a)** Expression of S490A mutation in full length *OCLN* or **(b)** S490A mutation in isoform 4 *OCLN*, inhibits VEGF driven proliferation as does **(c)** loss of extracellular loop 1 of *OCLN*. **(d-e)** BREC were transfected with GFP, WT or S490A *OCLN* and GFP-ΔNL1. **(d)** Cell cycle analysis was performed at 8 hours after VEGF treatment. Quantification from three independent experiments, each experiment consists of the cells from 2 individual wells in a 24-well plate, n=individual cell examined. Results are expressed as the percentage in each group with Chi-Square analysis. Expressing delta-loop 1 of *OCLN* altered cell cycle. **(e)** Detection of apoptotic cells by Click-iT TUNEL assay was measured at 8 hours after VEGF treatment. Quantification from three independent experiments, each experiment consists of the cells from 2 individual wells in a 24-well plate, n=individual cell examined. Results are expressed as the percentage in each group with Chi-Square analysis. **(f)** VEGF-induced permeability was measured in BREC transfected with empty vector, S490A in full length *OCLN* (FL-S490A) or S490A in *Iso4* (Iso-S490A) and analyzed by ANOVA and Sidak post-test.

### S490A mutation reduced OCLN dynamics

Previous studies revealed that OCLN-containing vesicles move along microtubes (MTs) ^35,36^ and together with the observed centrosomal localization, suggested that OCLN may traffic towards the minus-end of microtubules on dynein. Immunofluorescence staining of BREC with the antibodies against dynein, OCLN pS490, and α-tubulin demonstrated the co-localization of dynein and pS490 OCLN along microtubules. When cells were treated with Dynapyrazole to stall the dynein motor protein, the amount of co-localization was even greater (Fig. 4a). In addition, live cell trafficking experiments were carried out in U2OS cells transfected with GFP tagged *OCLN*WT or S490A or S490D mutants using spinning disk microscopy (Fig. 4b). Vesicles that ran longer than 5 seconds with displacement above 0.5 µm were evaluated for distance, displacement and speed. The results indicated that S490A mutation had modest but significant reduction in distance and displacement compared to wild-type *OCLN* and S490D mutants. (Fig. 4c). However, the largest impact was observed on the percentage of vesicles with displacement above 0.5 µm that was significantly reduced in GFP-S490A transfected cells compared to GFP-WT and GFP-S490D (Fig. 4c and d). These studies demonstrate colocalization of OCLN and dynein and reveal that S490A mutant reduces OCLN trafficking.

**Figure 4.**
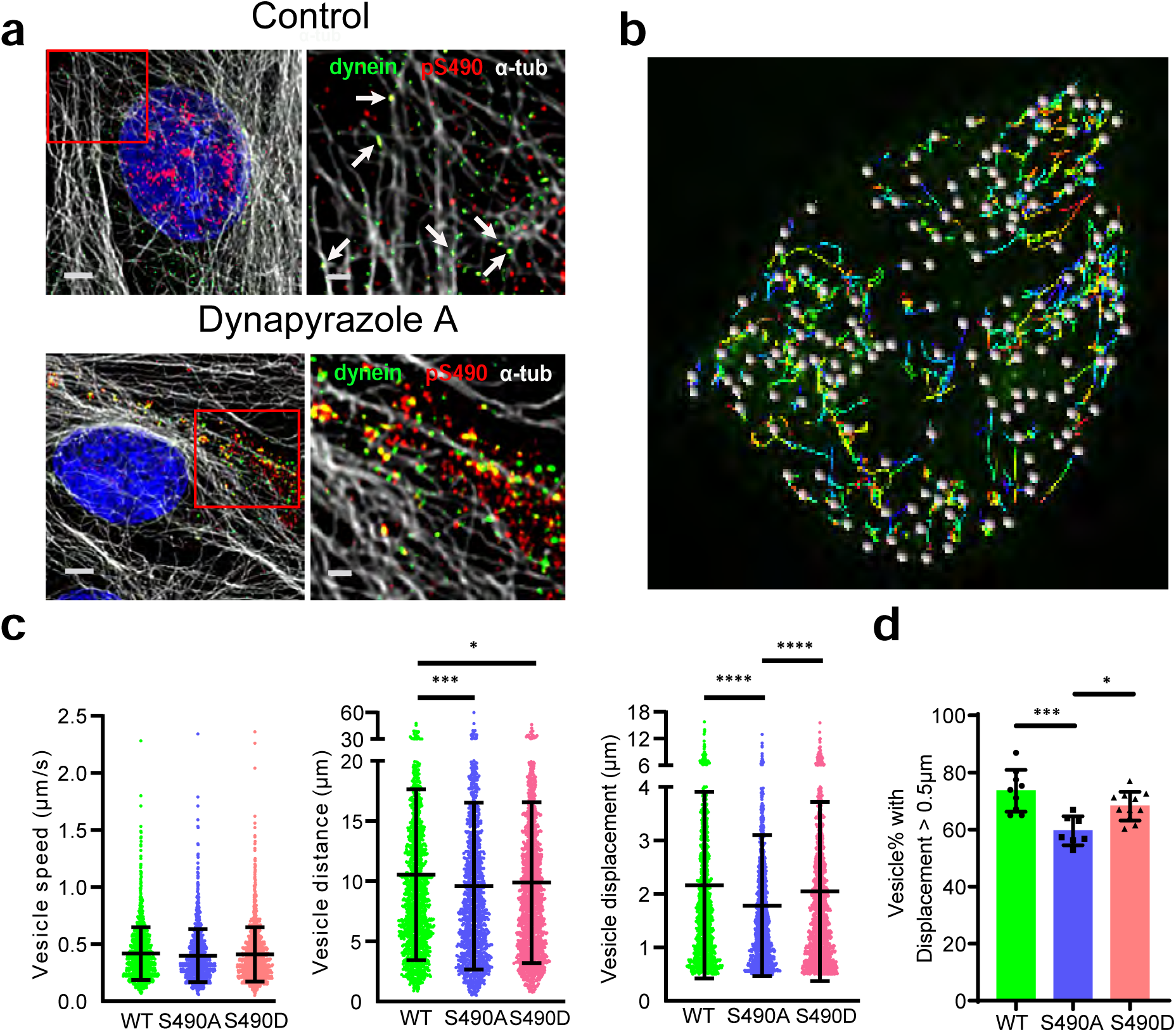
S490A mutation reduced *OCLN* dynamics. **(a)** Immunofluorescence (IF) staining with occludin pS490 (red), dynein (green), and α-Tubulin (white) revealed that OCLN traffics with dynein on microtubules in BREC. Top panel, treated with vehicle DMSO. The arrows shown in right zoomed-in panel indicate the colocalization of pS490, dynein, and α-Tubulin. Dynein trafficking was stalled in BREC treated with 10µM of Dynapyrazole A for 6h (bottom panels). Both pS490 and dynein were heavily accumulated and co-localized on microtubules, shown in the right zoomed-in panel. Scale bars: left panels, 3µm; right zoomed-in panels, 1µm. **(b)** Representative still images showing wildtype GFP-*OCLN* transfected U2OS cells during live cell imaging experiment and the moving tracks identified by Imaris software for quantification. **(c)** Quantification of vesicle speed, distance, displacement with each data point representing a vesicle with greater than 0.5µm displacement. **(d)** Percent of vesicles with greater than 0.5µm displacement per video from 2-4 transfections with each data point representing the data from a video.

### OCLN interacts with the LIC of the motor protein dynein

To explore the role of dynein in regulation of OCLN trafficking, we performed co-immunoprecipitation (co-IP) studies in U2OS cells and primary bovine retinal endothelial cells (BREC). IP of dynein targeting the intermediate chain (IC) co-precipitated OCLN as detected by both anti-OCLN (C-terminus) and pS490 Abs in U2OS cells (Fig. 5a) and in both confluent and nocodazole arrested, mitotic BREC cells (Fig 5d). Since many cargo adaptors specifically interact with the dynein light intermediate chain (LIC) 1 or 2 of dynein ^37–40^, we assessed occludin co-IP for this specific subunit of dynein. An OCLN IP led to co-precipitation of both dynein light intermediate chain 2 (LIC2) (Fig. 5b and e) and dynein light intermediate chain 1 (LIC1) (Fig. 5c).

**Figure 5.**
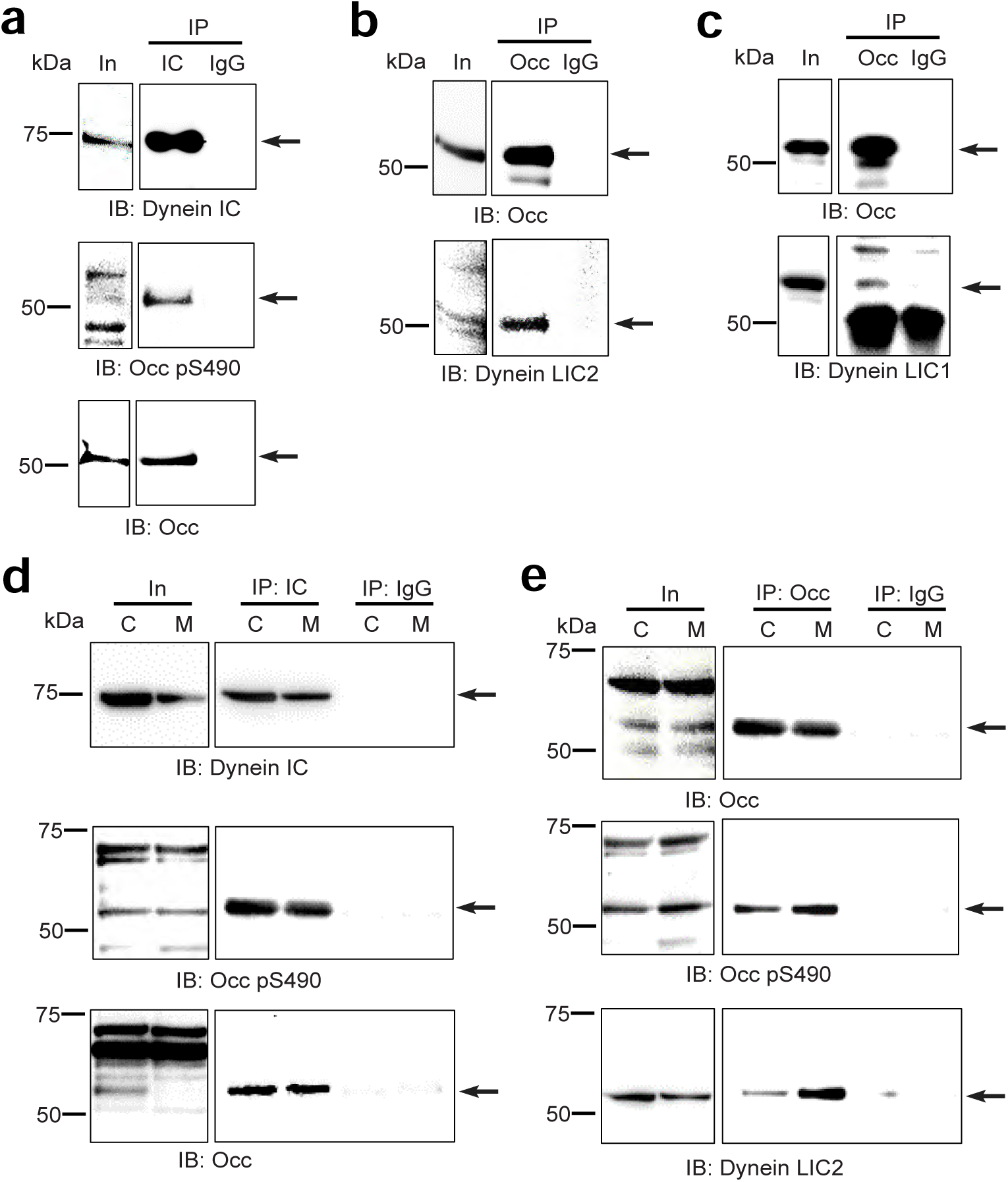
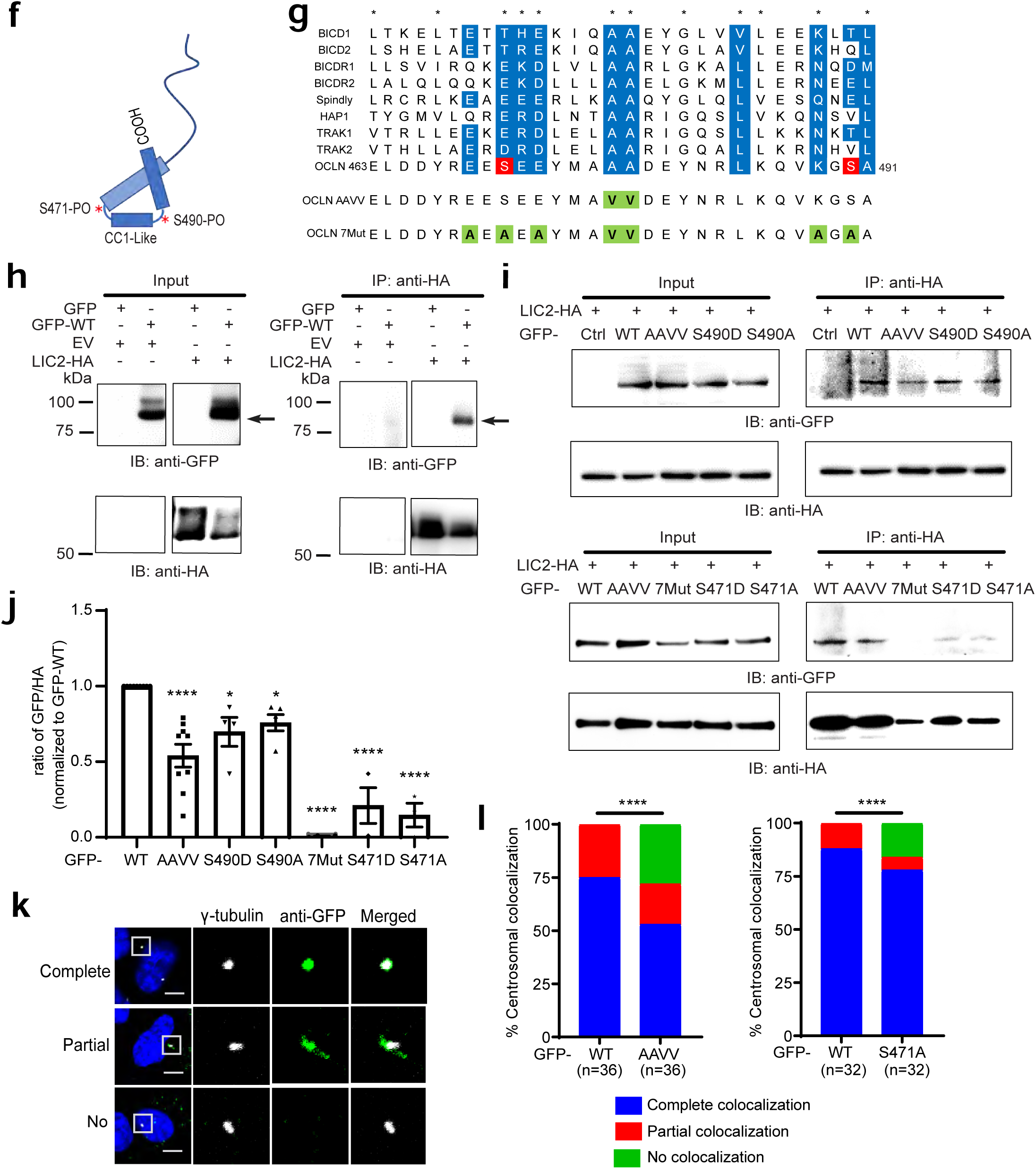
Occludin interacts with motor protein dynein LIC2. Immunoprecipitation (IP) of U2OS cell lysates with **(a)** anti-dynein IC or control IgG antibody (Ab). The input lysates (In) and IP elutes were immunoblotted (IB) with anti-dynein, anti-OCLN pS490 and anti-OCLN C-terminal Ab. **(b)** IP with anti-OCLN C-terminal or IgG Ab and IB with anti-OCLN, or anti-dynein LIC2. **(c)** IP with anti-OCLN C-terminal or IgG Ab and IB with mouse anti-OCLN, or rabbit anti-dynein LIC1 Ab. **(d** and **e)** IP from confluent (C) or mitotic (M) BREC. Mitotic BREC were treated with VEGF (50ng/ml) at 70% confluence for 30 min then synchronized with nocodazole (200ng/ml) for 16 hrs. **(d)** IP with mouse anti-dynein IC or IgG Ab and IB with mouse anti-dynein IC, OCLN pS490, and OCLN C-terminal Ab. **(e)** IP with anti-OCLN C-terminal or IgG Ab and IB with anti-OCLN, anti-OCLN pS490, and anti-dynein LIC2 Ab. **(f)** Schematic diagram of OCLN Isoform 4 with CC1-like box containing both S471 and S490 sites located in OCLN coiled-coil region. **(g)** Sequence alignment showing coiled-coil region of OCLN has homology with known dynein activators. Asterix represents known CC1-box homology, blue regions reveal consensus with OCLN. OCLN mutants highlighted in green. **(h)** U2OS cells co-transfected with *LIC2*-HA and GFP tagged wildtype *OCLN* (GFP-WT) and IP with rat anti-HA mAb. The input lysates and IP elutes were immunoblotted with rabbit anti-GFP pAb and rat anti-HA mAb. Capture assay reveals LIC2-HA co-precipitates GFP-WT occludin only when both genes are expressed. **(i)** U2OS cells co-transfected with *LIC2*-HA and GFP-WT or its mutants and IP with rat anti-HA mAb. The input lysates and IP elutes were IB with rabbit anti-GFP pAb and rat anti-HA mAb. **(j)** Quantification from at least three independent experiments. **(k)** Representative confocal images of U2OS cells transfected with GFP-tagged *OCLN* showing partial or no co-localization of GFP-OCLN and γ-tubulin as compared to complete colocalization group. Scale bars: 5 µm. **(l)** Quantification of γ-tubulin and OCLN colocalization as the percentage of each group by Chi-Square analysis.

There are four canonical dynein adaptor families, which are defined by the domain with which they bind to the LIC. Two of the most well studied adaptor families include those with a HOOK domain, for example HOOK microtubule tethering protein and those with a coiled coil 1 (CC1)-box, highlighted in bicaudal D homologue 2 (BICD2). The CC1 family interacts with the LIC via an AAXXG motif (where x denotes any amino acid) in the CC1-box. Mutations of this region can disrupt binding with LIC ^39^. Sequence alignment of OCLN and known dynein CC1-box containing adaptors (adapted from ^39^) revealed homology in OCLN CC domain that we referred to as CC1-like box. OCLN possesses an AAXXY rather than AAXXG motif plus additional novel sites of homology to the previously identified consensus. In all, 8 of 12 CC1-box consensus sites were conserved in OCLN and two additional sites of conservation were noted (Fig. 5f and g). Strikingly, these were found to include the Ser471 and S490 OCLN phosphorylation sites. To determine which region of OCLN interacts with dynein, capture assays were performed in U2OS cells that were transfected with GFP-tagged *OCLN* mutants and HA-tagged *LIC2*. As indicated in Fig.5h, LIC2-HA pulldown clearly co-precipitated WT-OCLN. Mutation of the 2 conserved alanine residues (A477 and A478) to valine (GFP-AAVV) decreased the LIC2 capture of OCLN by 50% (Fig. 5i and j). Both S490D and S490A mutations had only a modest 25% reduction in LIC2 capture. However, when we mutated 7 conserved sites of the OCLN CC1-like box (the two central Ala to Val (A477V, A478V), S471 and S490 (S471A and S490A) and 3 charged amino acids (E469A, E473A, K488A) a complete loss of binding to LIC2 was observed (Fig. 5i and j, Supplementary Fig. 2a and 2b). Further, more than 80% reduction of OCLN co-IP in LIC2 capture was found in both S471A and S471D mutants using an LIC2-HA capture (5j) or a LIC2-flag capture (Supplementary Fig. 2b) suggesting phosphorylation of this site per se was needed for complex formation with LIC.

We then tested how these mutations altered trafficking to centrosomes. GFP-OCLN co-localization with γ-tubulin was determined in U2OS cells as above. However, we noted partial co-localization and catalogued this as a separate group. The AAVV mutant reduced co-localization by 50% while S471A reduced co-localization by 25% (Fig. 5k and l). Consistently, the S471D mutant reduced co-localization by 40% while S490D reduced co-localization by only 20% (Supplementary Fig. 2c). Notably, none of these mutations were as robust in blocking centrosomal co-localization as was S490A suggesting the AAVV and the S471A mutant fail to act in a dominant manner like the S490A mutant.

We developed additional capture studies to test whether the coiled coil was necessary or sufficient for binding. U2OS cells were transfected with OCLN GFP-CTer mutants and HA-tagged LIC2. As indicated in Fig.6a, LIC2-HA successfully co-precipitated OCLN C-Ter as well as full length OCLN. However, LIC2-HA failed to co-precipitate occludin CC alone (Supplementary Fig. 3). Moreover, the binding to LIC2-HA was reduced by adding a glutathione S-transferase (GST) tag at the N-terminus of OCLN C-terminal (GFP-GST-Cter) to promote dimerization. Further, LIC2-HA pulldown still co-precipitated OCLN lacking the CC domain (GFP-ΔCC) (Fig 6b). These studies suggest the carboxy terminus has a second binding site for complex formation with LIC that does not require the coiled-coil domain. However, complex formation with endogenous OCLN cannot be ruled out.

**Figure 6.**
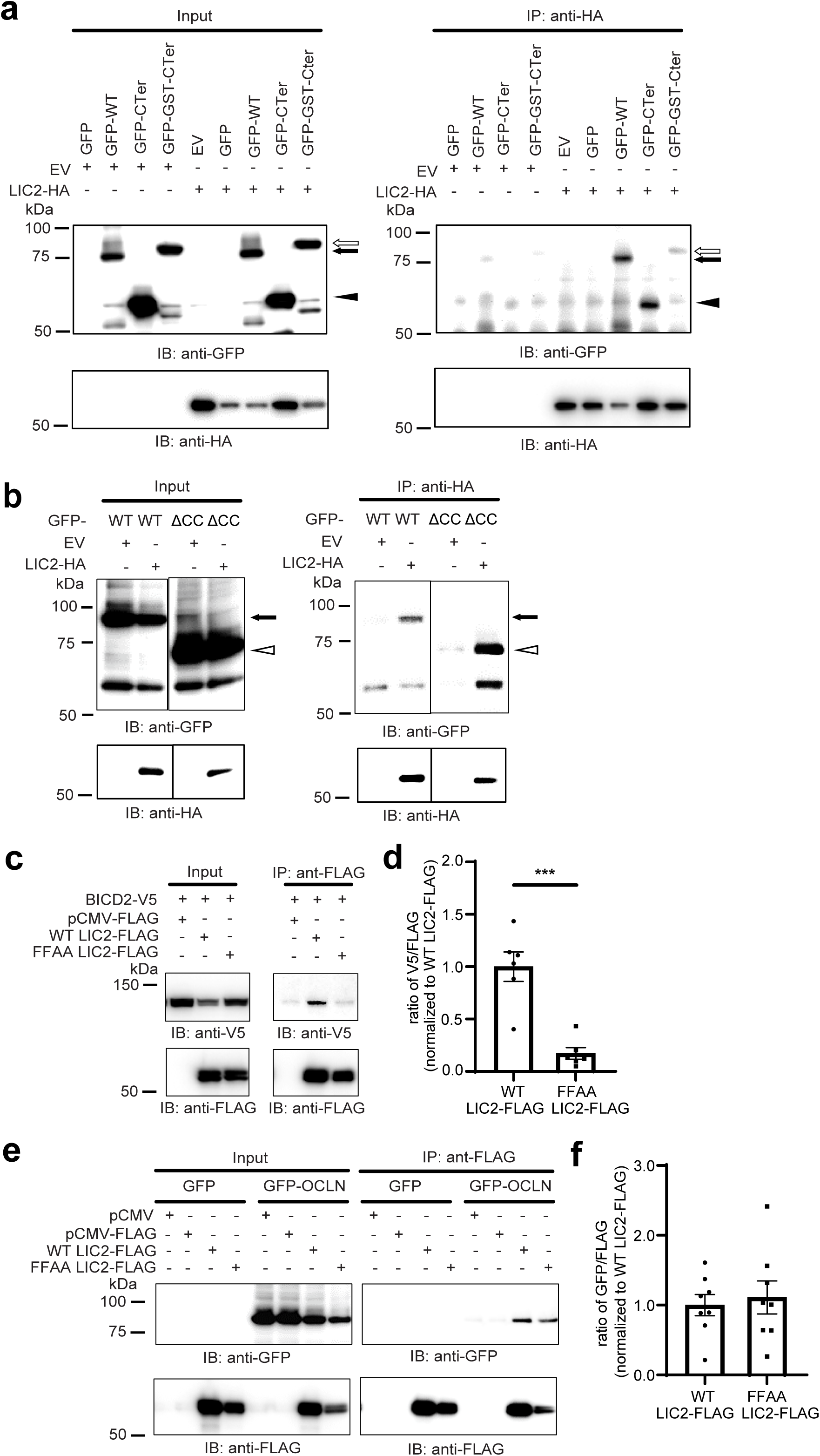
OCLN interacts with LIC2 through the coiled-coil domain and a separate binding site(s). **(a)** U2OS cells were co-transfected with *LIC2*-HA and N-terminal GFP tagged wild type *OCLN* (GFP-WT, arrow), GFP C-terminal OCLN (GFP-C-Ter, arrowhead), or GST forced dimerized C-terminal OCLN (GFP-GST-CTer, empty arrow) and IP with rat anti-HA mAb. The input lysates and IP elutes were IB with anti-GFP, anti-HA, and mouse anti-β-actin Ab. **(b)** U2OS cells were co-transfected with *LIC2*-HA and N-terminal GFP tagged wild type OCLN (GFP-WT, arrow) or GFP-WT with coiled-coil region deleted (GFP-Δ*CC*, empty arrowhead) and IP with rat anti-HA mAb. The input lysates and IP elutes were IB with anti-GFP Ab, anti-HA Ab. **(c)** U2OS cells were co-transfected with 3xFLAG tagged dynein *LIC2* (WT *LIC2*-FLAG), or *LIC2* mutant F432A/F433A (FFAA *LIC2*-FLAG), or control vector (pCMV-FLAG) and co-transfected with pcDNA5-StrepII-Halo-*BicD2*-V5-FRB (BiCD2-V5). 3xFLAG tagged LIC2 was captured with anti-FLAG M2 Ab and protein G Sepharose beads then eluted by 3xFLAG peptide. The input lysates and IP elutes were IB with anti-V5 and anti-FLAG M2 Abs. **(d)** Quantification from six independent experiments. **(e)** U2OS cells were co-transfected with WT *LIC2*-FLAG, FFAA *LIC2*-FLAG, or pCMV-FLAG and co-transfected with GFP or GFP tagged wild type *OCLN* (GFP-OCLN) for 48 hours. Cell lysates were IP for 3xFLAG tagged LIC2 with Protein G Sepharose beads/anti-FLAG M2 mAb and eluted by 3xFLAG peptide. The input lysates and IP elutes were immunoblotted with anti-GFP, anti-FLAG M2 mAb. **(f)** Quantification from eight independent experiments.

The studies suggested that the dynein LIC interaction with OCLN may be different as compared to the other known dynein adaptors. To test this hypothesis, we mutated the conserved hydrophobic residues F447 and F448 to alanine within LIC2 (FFAA LIC2) that was previously reported to significantly reduce LIC1 and BICD2 interaction ^39^. In agreement with the previous study, FFAA LIC2-FLAG failed to co-precipitate BICD2 as compared to its WT LIC2-FLAG control (Fig. 6c and d). In contrast, no significant difference was found between OCLN binding to FFAA LIC2-FLAG or its WT LIC2-FLAG control, indicating a distinct binding mechanism (Fig. 6e and f). Collectively, these data provide compelling evidence that OCLN interacts with LIC2 by the CC1-like box, and a second site in the carboxy terminus. Whether this represents direct interaction or a larger complex remains undetermined at this point.

### Homozygous deletion of full length and Iso4 *Ocln* is embryonic lethal and leads to blood vessel ruptures

The data indicates that OCLN carboxy terminus interacts with dynein through the LIC allowing trafficking in a phosphorylation dependent manner. Previous gene deletion targeting exon 3 failed to remove Iso4 *Ocln* ^25^. Therefore, we created a new mouse targeting exon 5 of *Ocln* to delete both full length and Iso4 *Ocln*. Mice were created in the University of Michigan transgenic core by CRISPR Cas9 targeted insertion of a gene cassette carrying LoxP recombination sites flanking exon 5 (*Ocln*^fl5/fl5^) and were sequence verified. *Ocln*^fl5/fl5^ mice were crossed with mice carrying Ella Cre to induce germline deletion of full length and Iso4 OCLN (Fig. 7a). Reduction of full length 55kDa, and modified 70kDa OCLN was confirmed by western blot analysis using C-terminal (Fig. 7b) and N-terminal (Fig. 7c) targeted antibodies, consistent with partial penetrance of Ella Cre ^41^. Reduction of Iso4 was observed with the C-terminal Ab only, as expected. Germline deletion of *Ocln*^fl5/fl5^ was found to be embryonic lethal (Fig. 7d). An analysis of embryos revealed normal Mendelian ratios at E14.5. However, by E18.5 the number of homozygous knock out animals began to decrease and no Cre^+^ *Occ*^fl5/fl5^ pups were observed at birth. Analysis of the embryos was performed by UM Pathology Core with two trained veterinarian pathologists. Random events of blood vessel rupture leading to hemorrhaging was observed in Cre^+^ *Occ*^fl5/fl5^ embryos examined. These ruptures occurred in the liver, brain and umbilical vein and artery (Fig. 7e-g). In addition, extensive blood accumulation was found in the pericardium. These ruptured vessels were considered the most likely cause of death. Two animals were observed with gastroschisis, with failure of proper midline closure and intestine exiting the cavity (Fig. 7h). The distinct difference from deletion of exon 5 that targets both full length and Iso4 *OCLN* and deletion of exon 3 targeting only full length *OCLN* highlights the essential role of the carboxy-terminus/Iso4 in development.

**Figure 7.**
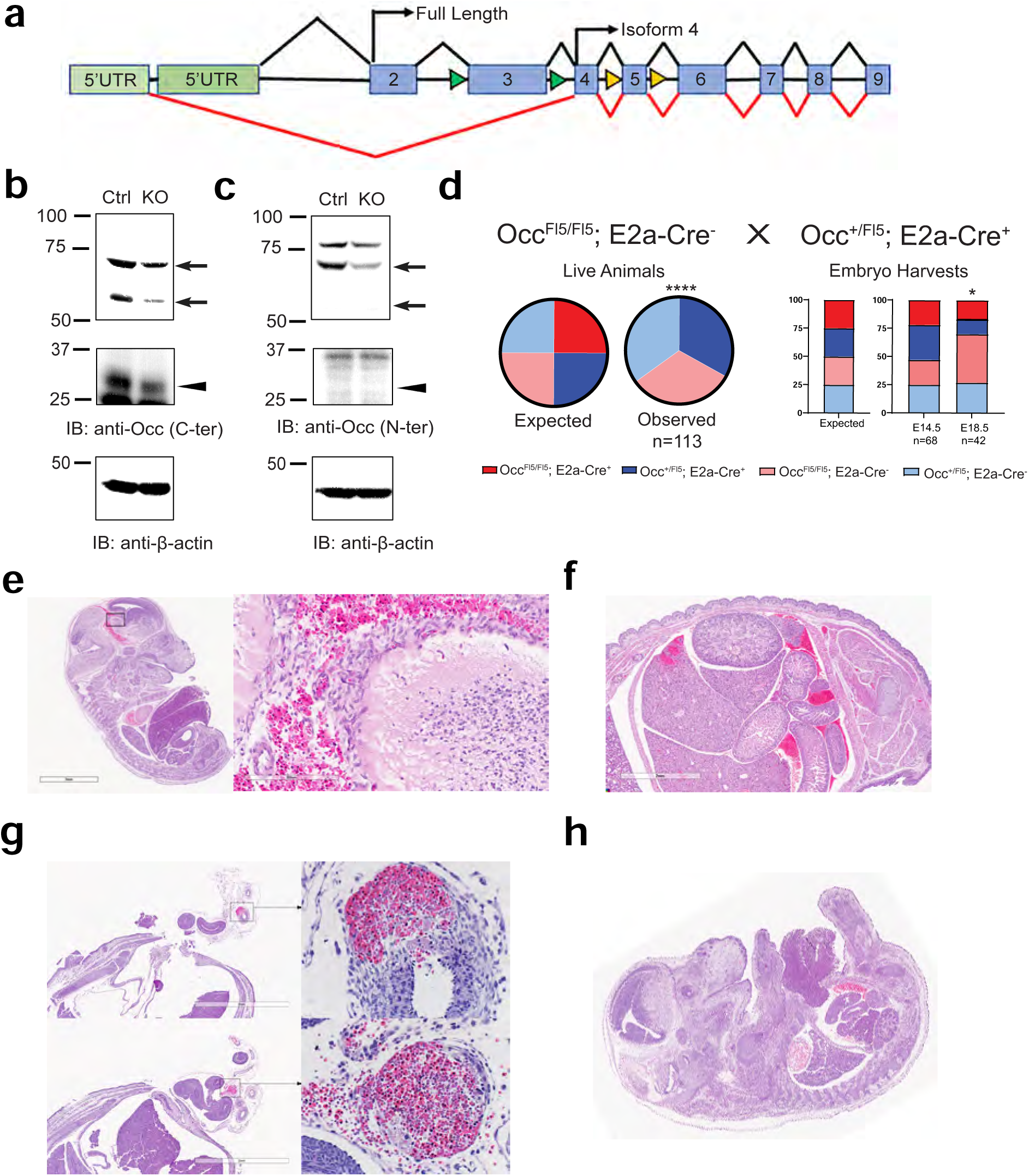
Homozygous deletion of *OCLN* from flex5 is embryonic lethal. (a) *OCLN* gene schematic showing full length and isoform 4 start sites with exon 3 floxed (fl3, green arrows) or the newly created exon 5 floxed (fl5, yellow arrows) mice. *Ocln*^fl5/fl5^ mice were crossed with germline Cre (E2A-Cre), *Ocln*^fl5/+^ mice to induce deletion of FL and Iso4 *Ocln*. **(b** and **c)** Protein was extracted from E15.5 whole embryos of Cre^-^ (Ctrl) or Cre^+^ *Ocln*^fl5/fl5^ mice followed by IB with **(b)** anti-C-terminal OCLN Ab, **(c)** anti-N-terminal OCLN Ab, or anti-β-actin Ab. Known location of full-length and isoform 4 mouse OCLN are indicated with the arrows and arrowheads respectively. **(d)** No Cre^+^, *Ocln*^fl5/fl5^ pups were observed at birth. An analysis of embryos revealed normal Mendelian ratios at E14.5. However, by E18.5 the number of homozygous knockout animals began to decrease. Quantification of the percentage of expected mendelian genetics and observed genetics was performed with Chi-Square analysis. **(e-g)** Analysis of the embryos reveals random events of blood vessel rupture leading to blood accumulation. These ruptured vessels were considered a likely cause of death. **(e)** H&E image showing cranial hemorrhage in E14.5 flex5 KO (sagittal). **(f)** H&E image showing liver/abdominal hemorrhage in E18.5 flex5 KO (coronal). **(g)** H&E image showing artery (top) and umbilical vein (bottom) hemorrhage in E14.5 flex5 KO (coronal). **(h)** H&E image showing gastroschisis in E14.5 flex5 KO (sagittal).

## Discussion

The current studies reveal that the OCLN carboxy terminus forms a complex with dynein through LIC. Mutational analysis suggests that the previously identified S471 phosphorylation site ^4,12^ is necessary for binding to LIC while S490 phosphorylation was necessary for trafficking as S490A mutations blocked centrosomal localization and reduced live cell trafficking of OCLN. Expression of a construct mimicking Iso4 splice variant of OCLN with the S490A mutation, blocked VEGF-induced cell proliferation and vascular permeability. Importantly, a novel gene deletion of OCLN targeting exon 5 that deleted both full length and Iso4 *OCLN*, was embryonic lethal. Previous studies targeting exon 3 were not lethal, revealing the requirement for the carboxy-terminus of OCLN in development. Expression of S490A mice appear to allow normal development but restrict VEGF induced permeability ^16,17^ and greatly impairs collateral vessel angiogenesis in the brain and the retina as shown here. Collectively, we propose that the OCLN carboxy terminus acts to link a vesicle of tight junction cargo to the dynein motor to allow trafficking from the cell border to the interior, leading to TJ strand breaks and vascular permeability. Further, OCLN trafficking to the centrosome appears essential for VEGF driven proliferation and collateral vessel formation (Fig. 8).

**Figure 8.**
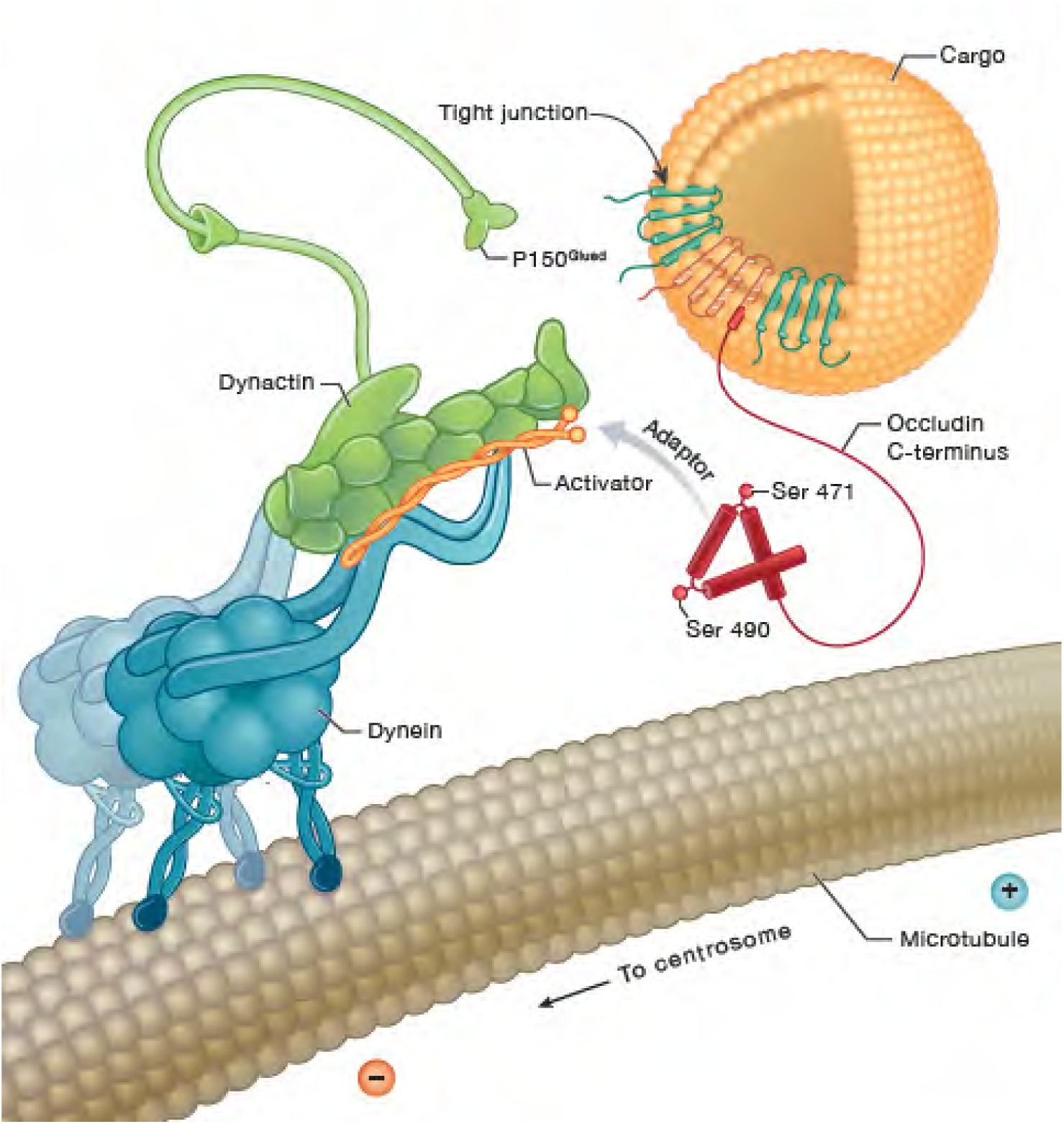
Occludin interacts with dynein to regulate both VEGF-induced permeability and angiogenesis. OCLN carboxy terminus interacts with LIC of dynein to allow trafficking of tight junction proteins as vesicular cargo from the cell border to the cell interior in response to VEGF leading to strand breaks and endothelial permeability in a Ser490 phosphorylation regulated manner. OCLN may further traffic to centrosomes in a vesicle necessary for endothelial proliferation and collateral vessel growth *in vivo*. Deletion of OCLN, including isoform 4, leads to embryonic lethality, demonstrating the importance of the OCLN carboxy terminus.

Previous studies have explored tight junction protein trafficking. Early studies revealed PDGF induced endocytosis and TJ strand breaks in Madine Darby canine kidney (MDCK) cells ^42^. OCLN was found to traffic on microtubules as observed by live cell imaging and dynein inhibitors led to accumulation of OCLN in the cytoplasm suggesting a role for dynein trafficking of OCLN to the plasma membrane ^35^. Also, in epithelial cells, Rab13 and junctional Rab13-binding protein (JRAB), also called molecule interacting with CasL like 2 (MICAL-L2), was required for OCLN and other tight junction and adherens junction protein recycling and barrier formation ^43,44^. Studies also demonstrated junctional trafficking in endothelial cells in response to VEGF that was dependent on OCLN S490 phosphorylation downstream of PKCβ ^14,15^. These studies demonstrated that OCLN may be ubiquitinated downstream of PKCβ phosphorylation and proteins with a ubiquitin interacting motif could chaperon OCLN. However, no evidence was found for ubiquitination of OCLN in co-precipitates with LIC and the Iso4 form of OCLN is lacking the N-terminal cytoplasmic region that interacts with the E3 ligase ITCH ^45^. It is common for transmembrane proteins to possess both non-ubiquitin and ubiquitin dependent trafficking pathways and OCLN appears to traffic in both pathways.

Dynein motor proteins drive minus end directed transport on microtubules. Cytoplasmic dynein-1, which is the only dynein motor that functions to facilitate cytoplasmic trafficking, is a 1.4MDa complex made of a heavy chain, intermediate chain, light intermediate chain (LIC) and 3 different light chains. All subunits of dynein are present as dimers in a motor ^40^. There are approximately 40 kinesin motors that primarily drive trafficking to the plus end of microtubules each with various cargo specificities, but dynein alone primarily drives minus end directed microtubule trafficking. Dynein achieves cargo specificity through specific adaptors that link cargo to dynein, in part by binding to the LIC ^40^. The canonical adaptors also act as bona-fide activators as demonstrated by *in vitro* processive trafficking using purified proteins. To activate dynein, adaptors form a complex with dynein and the co-factor dynactin. In this complex, dynein assumes a conformation that enables processive movement toward the minus end of microtubules and cargo transport ^37^. Activators generally possess a long coiled-coil domain with an LIC interaction site. Currently there are four established families. Two of the most well studied activator families are the Hook domain and the CC1-box domain containing binding motifs ^39^. Most activators possess a second spindly motif binding site, which mediates binding to dynactin ^46^. The CC1-box or Hook domain on the activator binds to an intrinsically disordered region of LIC that forms a transient helix upon binding thus allowing a range of activators to bind LIC ^38^.

Strikingly, OCLN possesses homology to the CC1-box within the C-terminal coiled coil domain and this region resides between the two turns containing the S471 and S490 phosphorylation sites. However, the canonical AAXXG motif was instead AAXXY in OCLN and thus we identify it as CC1-like box. Mutational analysis of this region revealed it is indeed necessary for full LIC binding. However, F447 and F448 mutated to alanine within LIC (FFAA LIC2) that abrogates binding to known canonical dynein adaptors ^38,47^, did not alter OCLN binding. Thus, OCLN with a CC1-like box may be more similar to the adaptor Rab7 interacting lysosomal protein (RILP) that binds LIC directly but also binds the homotypic fusion and protein sorting (HOPS) complex and is linked to dynein ^48^ or NuMA that has a hook domain and CC1-like box domain and interacts with LIC ^49^. The short coiled-coil domain of OCLN and the ability to bind LIC mutated at F447 and F448 suggests OCLN may not act directly to activate processive motility like established dynein activators ^40^. A second binding site allowing LIC complex formation was identified after expressing *OCLN* coiled coil deletion mutants and ongoing studies are underway to identify this site and assess direct or indirect interaction of OCLN with LIC.

Here, we establish that the coiled-coil domain of OCLN provides a phosphoregulatory role for LIC interaction and dynein trafficking. The strong effect of S471A or D on inhibiting binding suggests that phosphorylation specifically of this site is needed for OCLN binding to LIC that could not be substituted by the phosphomimic. Previous studies of this region revealed an interaction with ZO-1 also was impacted by phosphorylation at S471 and E470K or E472K had profound effect on ZO-1 binding ^4^. In all, this region binds to ZO-1 directly and contributes to LIC binding either directly or indirectly. Phosphorylation of S471 impacts binding to both ZO-1 and LIC. S490 phosphorylation was shown here to impact OCLN trafficking and S490A can act in a dominant negative manner. Clearly the possibility exists for competition of OCLN binding to ZO-1 or LIC for stable junction complex versus dynein trafficking which can be regulated by VEGF signal transduction.

The studies here are consistent with previous research revealing a role for the N-terminus of OCLN contributing to proliferation. Deletion of full length but not Iso4 *Ocln* from floxed exon 3 mice contributes to microcephaly and in organoids blocked neuroprogenitor proliferation with impaired spindle pole integrity ^25^ and human *OCLN* mutations lead to microcephaly ^29–31^. Here, mutations through the first extracellular loop (ΔNL1) led to reduced VEGF driven proliferation and increased apoptosis. However, S490A mutants completely blocked trafficking to centrosomes, prevented VEGF driven proliferation in cells but with limited apoptosis and blocked collateral angiogenesis in brain and retina strongly suggesting a required role for OCLN trafficking to centrosomes for endothelial proliferation. A recent study using exon 3 floxed mice revealed that loss of full length OCLN, but not Iso4, retarded mammary gland branching due to reduced cell proliferation associated with OCLN localization at centrosomes ^50^. Using biotin ID followed by capture assays for confirmation the investigators revealed OCLN interacts with the RAB11 family interacting protein 5 (FIP5) confirming previous biotin interaction studies identifying OCLN interaction with various endocytosis proteins including RAB11FIP1,2 and 5, RAB-5b, RAB-7a, RAB-10 and RAB-23 and vesicle associated membrane proteins 2 and 5 ^51^. The current and previous studies establish a role for the N-terminus of OCLN in cell proliferation and support OCLN vesicle trafficking leading to centrosomal localization.

Here we report for the first time deletion of both full length and Iso4 OCLN. Of note, deletion of full length and Iso4 OCLN with floxed exon 5 and a germline Cre led to embryonic lethality with vessel hemorrhaging as a likely cause of lethality. It was noted in the pathology report that points of hemorrhage appeared at regions of extensive branching. The cause for the midline opening and gastroschises is not known. However, gastroschisis is associated with improper development or rupture of the membrane covering the umbilical canal and umbilical ring ^52,53^. Here, multiple animals were observed with umbilical artery or vein rupture that could potentially lead to gastroschisis.

Other gene deletions also may lead to embryonic vessel hemorrhaging. Endothelial targeted gene deletion of β-catenin allowed normal vasculogenesis and angiogenesis but led to irregular vascular patterning and random vessel hemorrhaging observed at E10.5 ^54^. Further analysis revealed complete lack of capillary formation in the CNS with improperly formed endothelial aggregates and hemorrhage ^55^. Mutations in polycystin (PC or PKD)1 or 2 lead to autosomal dominant polycystic kidney disease ^56^. However, mutations of PKD1 also lead to life threatening vascular abnormalities and mouse embryos homozygous for PKD1 mutation (PKD1^L^) die by embryonic day 15.5 due to edema, vascular leaks and ruptured blood vessels ^57^. Future studies will use endothelial targeted Cre to elucidate the contribution of full length and Iso4 OCLN to vascular development and CNS barrier formation.

In conclusion, evidence here supports a role for the OCLN carboxy terminus to act as a non-canonical dynein adaptor linking vesicle cargo to this microtubule minus end directed motor through the LIC subunit. The interaction of OCLN to LIC depends on at least 2 regions with one a CC1-like box that exists in the OCLN coiled coil and is flanked by two previously identified phosphorylation sites, S471 found necessary for proper LIC binding and S490 required for regulation of trafficking. OCLN links tight junction containing vesicles to dynein leading to endocytosis and formation of gaps and increased permeability downstream of VEGF signaling while OCLN trafficking to centrosomes is necessary for proper VEGF driven endothelial collateral angiogenesis.

## Methods

### Reagents

The details of reagents used for the project are listed in the Supplementary table.

### Animals

All experiments conformed to the ARVO Statement for the Use of Animals in Ophthalmic and Vision Research and the guidelines established by the Institutional Animal Care & Use Committee (IACUC) of the University of Michigan. Transgenic mice were created with wildtype occludin (WT*OCLN*) or occludin containing Ser 490 mutated to Ala (*OCLN*S490A) under the CAG promoter followed by a floxed stop sequence to allow conditional expression ^16^ and were crossed with TEK-Cre mice from The Jackson Laboratory or *PDGFb*-iCreER mice received from Dr. Fruttiger ^58^ for tamoxifen inducible expression of S490A *OCLN* in endothelial cells. Mice with endogenous *Ocln* exon 3 floxed (*Ocln*^fl3/fl3^) that was obtained from Dr. Turner ^59^. *Ocln*^fl3/fl3^ mouse with Cre excision still allow expression of alternatively spliced isoform 4 of *Ocln* ^25^; therefore, mice with exon 5 of *Ocln* floxed (*Ocln*^fl5/fl5^) were generated for inhibiting production of full length occludin and isoform 4. *Ocln*^fl5/fl5^ mice were crossed with germinal Ella Cre (E2a Cre) mice from The Jackson Laboratory for germ line deletion of occludin.

### Histology analysis of mouse embryos

Mouse embryos were collected at ED14.5 or ED18.5 and fixed in 10% neutral buffered formalin at a ratio of 10:1 fixative: tissue for more than 24 hours before submitting the samples to the University of Michigan Unit for Laboratory Animal Medicine (ULAM) Pathology Core for histology analysis. Embryos were paraffin embedded using an automated tissue processor (TissueTek VIP5, Sakura). Tissues were sectioned on a rotary microtome at 4μm thickness at 50-200µm intervals (head 50µm, trunk 100µm, whole embryo 200µm) between each section. Slides were stained with H&E by standard methods (Autostainer ST5010 XL, Leica Biosystems). Light microscopic evaluation was performed at magnifications ranging from 20x to 600x. The evaluation was performed initially by a board-certified veterinary pathologist with peer review by a second board-certified veterinary pathologist. For images, representative slides were scanned on a digital slide scanner (Aperio AT2, Leica Biosystems) at magnifications up to 40x (0.25μm/pixel). Representative images were taken as Tiff files from the digitized slides using freely available software (Aperio ImageScope v 12.4.6.7001, Leica Biosystems).

### Branch retinal vein occlusion (BRVO)

BRVO was performed in control and transgenic mice (14–16 weeks) using a 532nm laser with a Micron III retinal imaging system (Phoenix Research Laboratories, Pleasanton, CA). Briefly, animals were anesthetized with intraperitoneal injection of Ketamine (100mg/kg, Hopira, Lake Forest, IL) and Xylazine (10mg/kg; Akorn, Lake Forest, IL). The pupils of anesthetized animals were dilated with topical 2.5% phenylephrine (Paragon BioTek, Inc., Portland, OR) and 0.5% tropicamide (Akorn, Lake Forest, IL). Fluorescein angiography (FA) was performed during the procedure and fluorescein fundus images were captured at 3– 5min after i.p. injection of 0.1ml of 2% fluorescein sodium (Akorn, Lake Forest, IL) before applying the laser spots. Laser spots were applied using a Meridian Merilas 532nm laser (Thun, Switzerland) connected to the Micron III with a laser injector (Phoenix Research Laboratories). The Micron III laser injector only delivers ∼7% of the energy to the eye, so the parameters were adjusted accordingly, and the optimized conditions used were: 2,000mW, 1000ms, and 50μm spot size. 8 to 12 laser burns were applied to a branch vein of each eye (2 to 3 disc diameters from optic disc). Fluorescein fundus images were re-captured after applying the laser spots for assuring complete blockage of blood flow of targeted retinal vein. On day 7 or day 28 after BRVO procedure, eyes were enucleated and fixed in 4% paraformaldehyde (PFA). Whole mount retinas were immunostained targeting CD31 and Ki67. To quantify cell proliferation and retinal neovascularization, a confocal Z-stack of 10 images were collected over a depth of 10 µm and projected as a single image. For each mouse, five fields surrounding the occluded vein was imaged and evaluated using AngioTool 0.6a ^60^ and averaged. Results represent 3 independent experiments.

Laser-Induced choroidal neovascularization (CNV) was performed in control and transgenic mice (6–8 weeks) using a Meridian Merilas 532 nm laser (Thun, Switzerland) connected to the Micron III with a laser injector (Phoenix Research Laboratories) as described previously ^61^.

### Mouse retinal explants

Retinal explants were maintained in 3D collagen matrix using a modification of protocol reported previously^62–65^. Briefly, a stock solution of rat type I collagen at 3mg/mL was diluted and mixed in 8:1:1 ratio with 10xPBS and 10X retinal explant medium prepared with endothelial basal medium containing 200mM HEPES, 5µg/mL fibronectin and laminin, 20mg/mL NaHCO3, and 0.2N NaOH. The mixed collagen solution was neutralized with 1N HCL to adjust pH to 7.4 to 7.5 and was kept at 37°C for 1h for polymerization. Meanwhile, the mouse eyes were enucleated immediately after euthanasia and kept in PBS. Each retina was dissected under a dissecting microscope and four relaxing cuts were made to obtain four evenly sized quadrants. Each quadrant was further cut into four 1x1mm pieces, yielding 16 pieces per eye for explants. Each piece was placed on the top of pre-gelled 400µl of 3D collagen matrix in a 24-well tissue culture plate. 150µl of the collagen gel mixture was added to each well and the plates were incubated at 37°C for 1h to allow polymerization of the upper gel layer. Lastly, the collagen gel sandwich was overlaid with 500µl of endothelial basal medium containing 10% horse serum (Lonza, Walkersville, MD), 12µg/ml bovine brain extract (Lonza, Walkersville, MD), 100µg/ml heparin, and 25ng/ml VEGF-A. Culture medium was replaced every 48h. On day 21, the samples were fixed with 4% of PFA and stained with IB4-Alexa 488 and Hoechst 33142 for overnight at 4°C. The stained samples were mounted onto glass slides with ProLong Gold mountant and coverslips. Outgrowth of vessels from explant samples were imaged on Leica DM6000 upright fluorescence microscope (Leica, Wetzlar, Germany) and evaluated using MetaMorph software version 7.6.3 (Molecular Devices, Sunnyvale, CA).

### Middle cerebral artery occlusion (MCAO)

Photothrombotic MCAO was induced using a 3.5-mW green laser (540 nm, Melles Griot) and Rose Bengal (RB) dye (Fisher) as described ^66^. Occlusion was achieved when the tissue perfusion unit dropped to less than 30% of pre-occlusion levels. To achieve a stable clot the laser was left on for 20 minutes. To evaluate post-stroke angiogenesis, mice were subjected to MCAO and 21 days later were anesthetized with isoflurane and euthanized by trans cardiac perfusion with PBS followed by 4% PFA. The brains were then post-fixed overnight in 4% PFA, transferred to 30% sucrose, and then cryopreserved in OCT prior to preparation of 16μm frozen sections. CD31 stained vessels were imaged at 4 regions next to the infarct of each section and three sections were analyzed for each mouse and quantified using AngioTool version 0.6a ^60^.

### Cerebral blood volume

Relative tissue blood volume in the infarct region in mice was measured after MCAO by MRI imaging as described previously ^67^. Briefly, T2-weighted images were recorded every 2 minutes starting before gadolinium contrast agent was administered (10mmol/kg, i.p.) and followed for 30 minutes after injection as the T2 signal saturates. The contrast agent was administered via a 26G Abbocath®-T vascular catheter (Hospira) that was inserted into the peritoneum cavity and connected to a catheter extension set (Infusion Devices) and 1mL syringe. The relative blood volume was calculated using the formula: BV∞ΔR2 = ln(Spre/Spost)/TE 72, where TE is the effective echo time, Spre is the T2-weighted signal before contrast agent administration and Spost is the T2-weighted signal after the contrast administration.

### Plasmid construction

Constructs used include: N-terminal GFP-tagged WT, S471A or S490A *OCLN* constructs in pmaxFP-Green-C expression vector (Lonza, Walkersville, MD) and also include mutants lacking N-terminal (*GFP-*Δ*N*) and extracellular loop 1(*GFP-*Δ*NL1*), *GFP-*Δ*NL1* mutant replacing loop2 with loop5 of Na^+^/H^+^ exchanger (*GFP-*Δ*NL1L2NHE*), or mutants for replacement of loop 1 with loop5 of Na^+^/H^+^ exchanger (*GFP-*Δ*NL1NHE*), mutants lacking C-terminal coiled-coil domain (*GFP-*Δ*CC*), C-terminal coiled-coil domain (*GFP-CC*), C-terminal (*GFP-C-Ter*). Other mutants include C-terminal V5-tagged *OCLN* in pmaxCloning expression vector (Amaxa Biosystems). Additional mutants were constructed mimicking the naturally occurring splice variant ^25,36^ starting at Met252 and containing 14 transmembrane amino acids and the cytoplasmic tail to the carboxy terminus of occludin (Iso4-V5). Also, an N-terminal myristylation signal (MGSSKSKPK) ^68^ was added to C-terminal *OCLN* (*Myr-CTer-V5*). BicD2-V5 construct was described previously ^69^. Mutants were made through the service provided by Genscript USA (Piscataway, NJ) and all constructs were confirmed by DNA sequencing.

### Cell culture, transfection, and immunocytochemistry

BREC were isolated and cultured as described previously ^14^. Transfections with plasmids were performed using Amaxa Nucleofector System (Amaxa, Koeln, Germany) for BREC and Lipofectamine 2000 (ThermoFisher Scientific, Waltham, MA) for U2OS cells. BREC and U2OS cells immunohistochemistry was performed on cells grown on either chambered cover glass (ThermoFisher MA) or iBidi dishes (iBidi USA) and fixed with methanol at 20°C for 10 min as previously described ^70^. After rehydration with PBS for 20 min, cells were blocked in 1% wt/vol BSA-PBS and incubated with primary antibodies overnight at 4°C, washed, and followed by a 1h incubation with secondary antibodies and Hoechst 33142 at room temperature. Primary antibodies were anti-turboGFP (OriGene Technologies, Rockville, MD), anti-V5 (Thermo Fisher. Scientific, Waltham, MA), anti-γ-tubulin (Millipore sigma, Burlington, MA), anti-OCLN and pS490OCLN previously characterized in ^12^. Additional information for the antibodies used for the study is provided in the Supplementary table. Secondary antibodies used were Alexa Fluor 488, 594, and 647 goat or donkey anti–rabbit, donkey anti-goat, donkey anti–mouse, and donkey anti–rat IgGs (Thermo Fisher or Jackson ImmunoResearch Laboratories, Inc.). Mice retinal flat mounts or brain cryostat sections were blocked in 10% donkey serum with 0.3% Triton X-100 for 1 h at room temperature. Samples were then incubated with primary antibodies and IB4 for 3 d at 4°C, followed with overnight incubation with secondary antibodies at 4°C. Samples were mounted with coverslips using ProLong Gold mountant. Samples were imaged using a TCS SP5 or Stellaris 8 confocal microscope (Leica, Wetzlar, Germany) and images were processed by Leica LAS AF software. Centrosomal co-localization of γ-tubulin and occludin or its mutants was quantified in a masked manner by three unbiased observers. Minimum 30 cells for each condition were analyzed.

### Cell proliferation and apoptosis Assays

BREC DNA synthesis and apoptosis assays were performed according to a modified protocol of Click-iT EdU Alexa Fluor 594 Imaging kit ^26^ and Click-iT Plus Alexa 647 TUNEL assay (Thermo Fisher). Briefly, BRECs were transfected with empty vector or occludin mutants and plated on pre-gelled 3D collagen matrix prepared as described above. After 24h, 150µl of the collagen gel mixture was added to each well and the plates were incubated at 37°C for 1h to allow polymerization of the upper gel layer. Lastly, the collagen gel sandwich was overlaid with 500µl of stepdown medium containing 1% fetal bovine serum and 50ng/ml VEGF-A. 10μM EdU was added to the cells at 6h for DNA synthesis study. At 8h, the samples were collected after dissolving the collagen gels with 1mg/mL type I collagenase and disassociating cells with 0.05% trypsin. The cells were further centrifuged at 500g for 5 min onto ThermanoxTM plastic round coverslips with a 13-mm diameter (Thermo Fisher) that were pre-coated with 1µg/cm^2^ of fibronectin bovine plasma (Millipore sigma, Burlington, MA) in 24-well plate for 1h at room temperature. The coverslips were mounted onto glass slides with ProLong™ Gold Antifade Mountant (Thermo Fisher). Cells samples were imaged using a fluorescent microscope (DM6000, Leica, Wetzlar, Germany) for DNA synthesis and apoptosis analysis, or a TCS SP5 or Stellaris 8 (Leica) confocal microscope for cell cycle analysis.

### In Vitro Permeability Assay

BREC permeability to 70-kDa RITC dextran (Sigma) after treatment with VEGF (50ng/ml) was determined by measuring the rate of flux of the dextran over a 4-h time course followed by calculating the diffusive permeability (Po) across the monolayer as described previously ^71,72^

### Live cell imaging

U2OS cells were transfected with *GFP-WT*, *GFP-S490A* or *GFP-S490D OCLN* and grown in the presence of neomycin (1.6 mg/ml) for 7 days before live cell imaging. GFP signal was imaged using a Yokogawa W1 confocal scan head mounted to a Nikon Ti2 microscope with an Apo TIRF 60 × 1.49 NA objective. The microscope was run with NIS Elements using the 488 nm line of the LUN-F-XL laser engine and Prime95B camera (Photometrics). The videos were recorded at 1frame/0.5s for 1min at single focal plane. Quantification of GFP speed, distance and displacement was performed by Imaris version 9.5.1 software with particle tracking algorithm. GFP signals that could be tracked for more than 5sec were evaluated.

### Capture Assay

Protein interactions were examined using capture assay according to a modified protocol described previously ^14^. Briefly, BREC or U2OS cells transfected with GFP tagged occludin and mutants, V5-tagged *BICD2*, HA-tagged or Flag-tagged dynein light intermediate chain 2 (*LIC2*) were lysed with co-immunoprecipitation (Co-IP) buffer (1% Nonidet P-40, 10% glycerol, 50mM Tris, pH 7.5, 150mM NaCl, 2mM EDTA, 1mM NaVO4, 10mM Na fluoride, 10mM Na-pyrophosphate, 1mM benzamidine, complete protease inhibitor and PhosSTOP tablet). The lysate was centrifuged at 12,000*g* for 10min, and the supernatant was transferred to another microcentrifuge tube. 500µg lysate was precleared using 60µl of Sepharose Protein G beads (50:50 slurry). Following clearing, 10µg of anti-occludin, anti-dynein LIC2, 0.1µg of anti-HA, or 3µg of anti-FLAG antibody was added to the lysate and incubated overnight at 4°C. The following day, lysates were incubated with 60µl of Sepharose Protein G beads (50:50slurry) for 1h at 4°C. Lysates were then centrifuged at 12000 x *g* for 1min. The supernatant was removed, and the pellet was washed with 500µl of Co-IP buffer for a minimum of 3 times. Co-IP buffer with 500mM NaCl was used for Co-IP of *GFP-*Δ*CC* or *GFP-CC* with *LIC2-HA*. Following the final wash, the pellet was resuspended in 60μL of low pH (2.8) glycine buffer for 5min at room temperature, then NuPage sample buffer (LDS/DTT mix) was added and samples were boiled for 10 minutes centrifuged at 12000 x g for 2 min and supernatant collected for immunoblotting analysis using the NuPAGE system (Thermo Fisher) as described previously ^26,73^. After blocking with milk in Tris-buffered saline with 0.1% Tween 20, immunoblotting was performed by incubating primary antibodies overnight at 4°C and goat anti-mouse, anti-rabbit, or anti-rat secondary antibody conjugated with horseradish peroxidase (1:10,000) or with alkaline phosphatase (1:10,000) and chemiluminescence with HRP substrate Lumigen TMA-6 (Lumigen, Southfield, MI) or AP substrate ECF (GE Healthcare, Piscataway, NJ).

### Statistical analysis

All experiments were repeated three or more times and expressed as the mean ± SEM and were analyzed using two-tailed student’s *t* test, ANOVA for three or more conditions with Sidak post-test, or Chi-Square analysis for frequency, using Prism 8.0 (GraphPad Software, La Jolla, CA) with: *P < 0.05, **P < 0.01, ***P<0.001 and ****P < 0.0001.

## Acknowledgements

We would like to thank Ingrid L. Bergin VMD, MS, DACLAM, DACVP and Yao Lee DVM, MS, PhD, DACVP of the University of Michigan, ULAM Pathology Core, RRID:SCR 018823. This research was supported by grants from the National Institutes of Health, EY012021 (DAA), HL055374 (DL, DAA), Vision Research Core Grant P30EY007003, the Michigan Diabetes Research and Training Center Grant P30DK020572, Research to Prevent Blindness (DAA) and NIH instrumentation grant S10OD028612.

## Author contributions

Concept and design: D.A.A., X.L., M.E.R., M.E.D., and D.A.L. Data Acquisition and analysis or interpretation of data: D.A.A., X.L., E.J.S., A.D., J.P.G., M.M., L.G., J.S.P., C.M.L., M.E.R., M.E.D., and D.A.L. Drafting of the manuscript: D.A.A. and X.L.

---

**Supplementary Figure 1.**
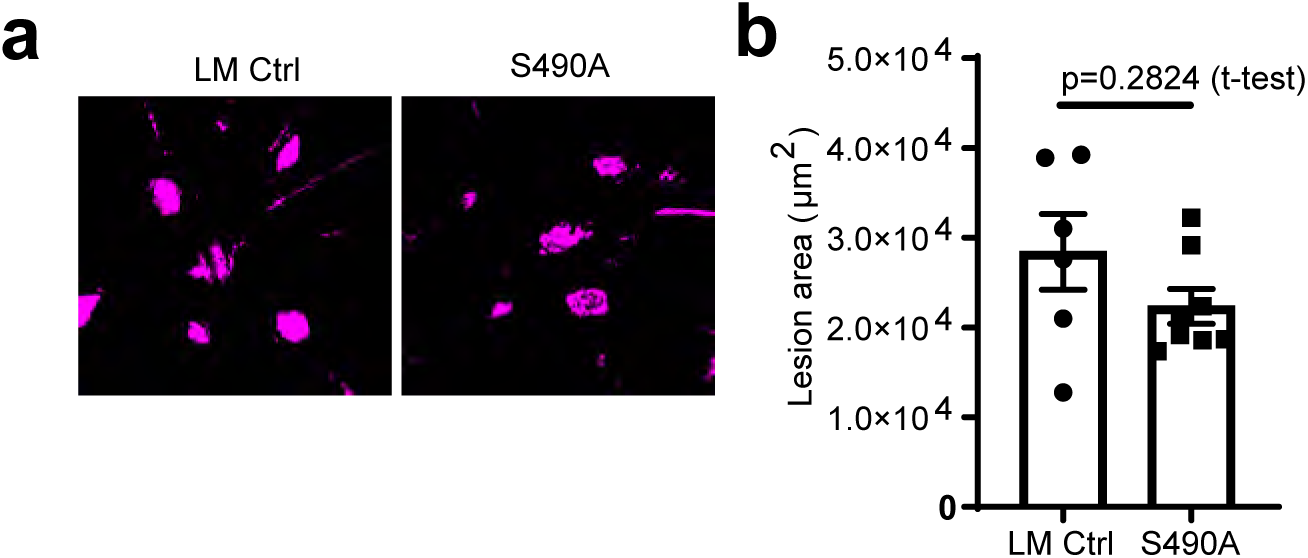
Expression of *OCLN* S490A and neovascularization in mice at 1 week after laser-induced choroidal neovascularization (CNV). **(a)** Representative epifluorescence images showing the laser-induced CNV lesions in the whole mount retinas stained with IB4 (purple). **(b)** Quantification of the lesion areas. Student t-test. P=0.2824. LM Ctrl: Tek-Cre^-^, *Ocln*^fl3/fl3^, *OCLN*S490A^+/+^; S490A: Tek-Cre^+^, *Ocln*^fl3/fl3^, *OCLN*S490A^+/+^.

**Supplementary Figure 2.**
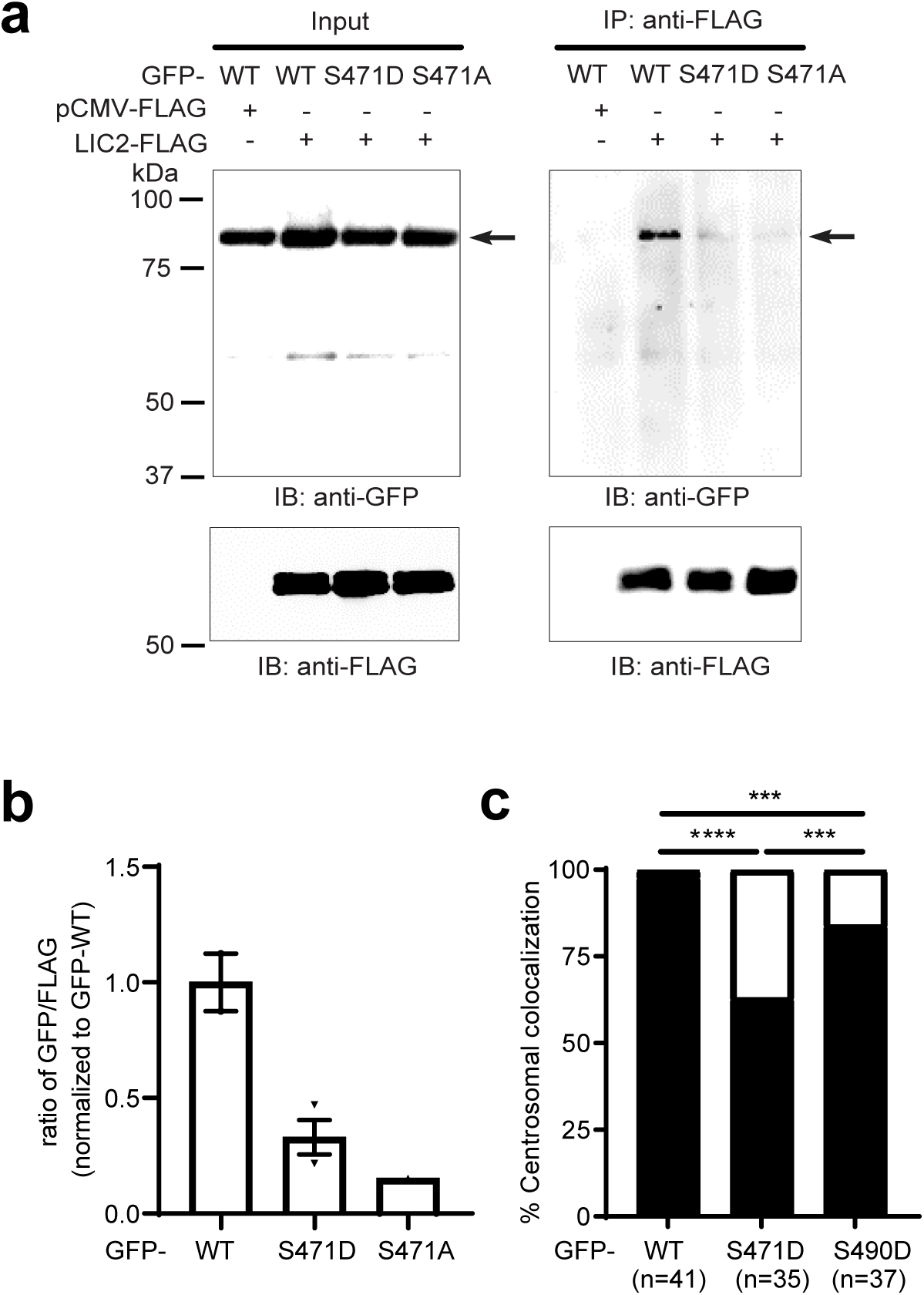
Expression of OCLN S471 mutants reduced occludin binding to dynein LIC2. **(a)** U2OS cells were co-transfected with *LIC2*-FLAG, or pCMV-FLAG with GFP-WT or S471 *OCLN* mutants (S471D or S471A) for 48 hours. 3xFLAG tagged LIC2 was captured with anti-FLAG M2 mAb and protein G Sepharose beads then eluted by 3xFLAG peptide. The input lysates and IP elutes were IB with goat anti-turboGFP, mouse anti-FLAG M2 Abs. **(b)** Results of the quantification from IP experiments are expressed as the mean relative to the WT OCLN ± S.E.M. **(c)** Quantification of the percentage of transfected U2OS cells with γ-tubulin and OCLN colocalization. Black bars, complete colocalization; open bars, no colocalization. Results are expressed as the percentage in each group with Chi-Square analysis. ***P<0.001 and ****P<0.0001.

**Supplementary Figure 3.**
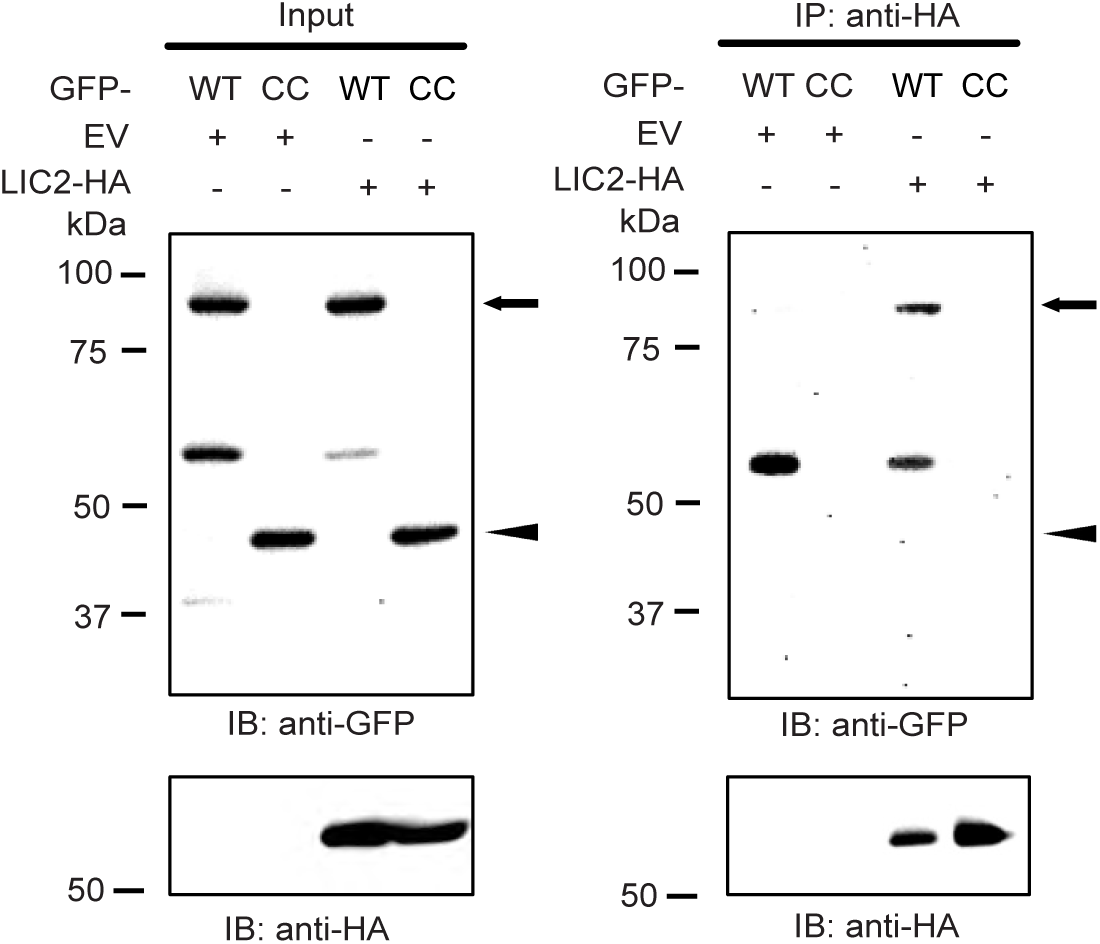
OCLN coiled-coil domain alone was not shown to interact with LIC2. U2OS cells were co-transfected with LIC2-HA and N-terminal GFP tagged wild type *OCLN* (GFP-WT, arrow) or GFP-coiled-coil region (GFP-CC, arrowhead) and IP with rat anti-HA mAb. The input lysates and IP elutes were IB with goat anti-GFP pAb, rat anti-HA mAb.

**Supplementary table 1.**
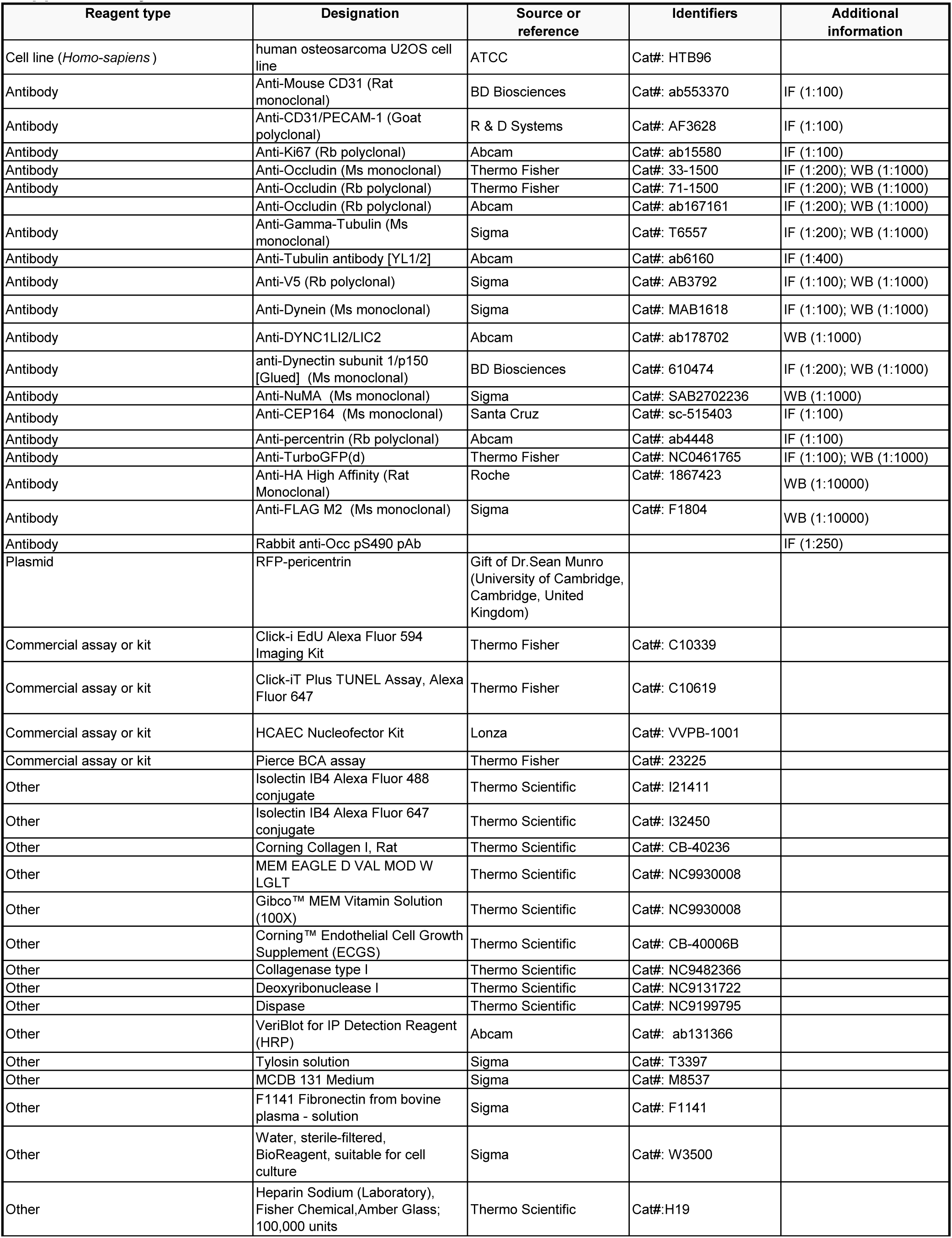
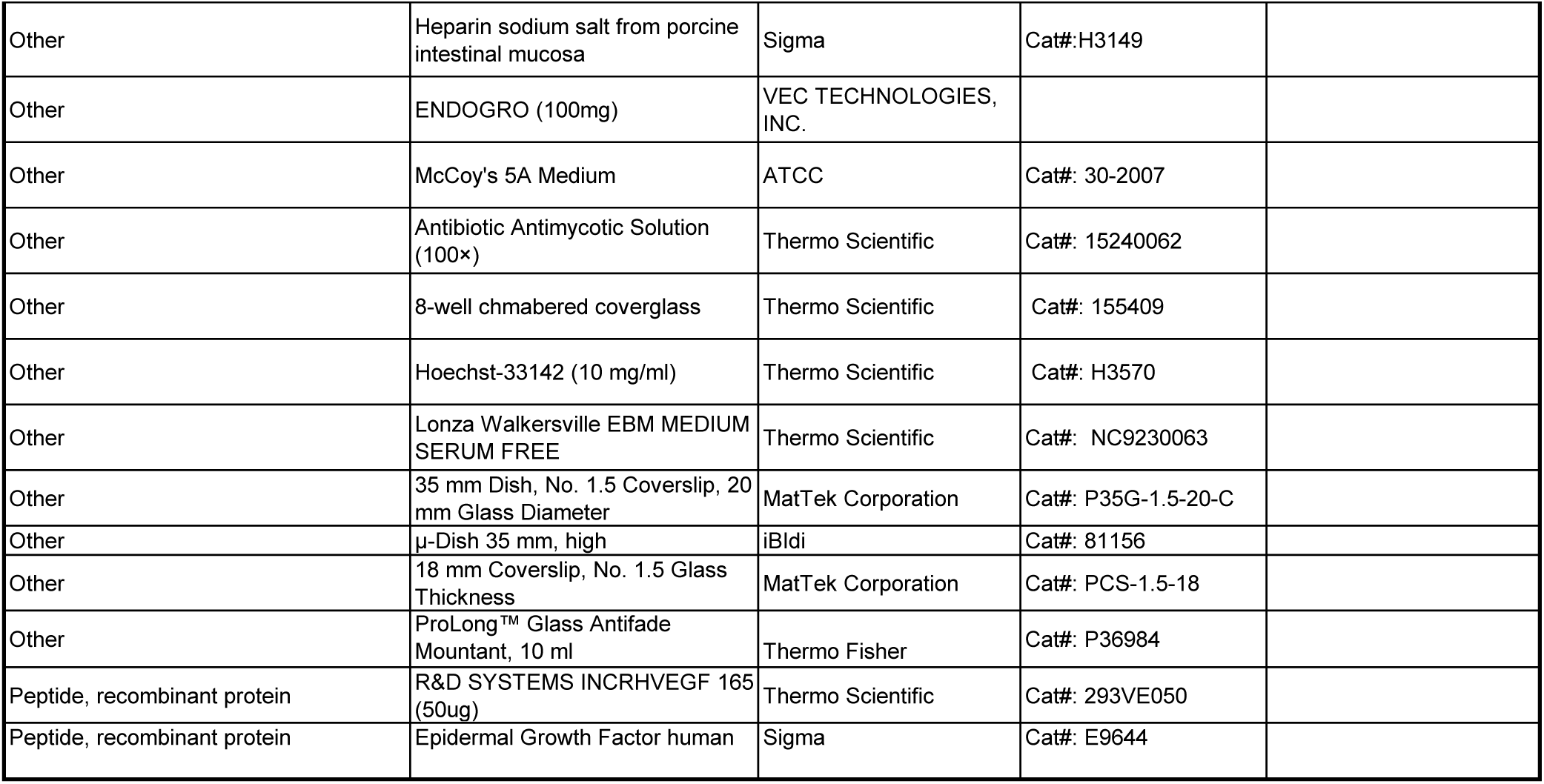

